# Signaling modality within gp130 receptor enhances tissue regeneration

**DOI:** 10.1101/2022.01.05.475124

**Authors:** Ruzanna Shkhyan, Candace Flynn, Emma Lamoure, Ben Van Handel, Arijita Sarkar, Jinxiu Li, Jesse York, Nicholas Banks, Robert Van der Horst, Nancy Q. Liu, Siyoung Lee, Paul Bajaj, Kanagasabai Vadivel, Hans I-Chen Harn, Thomas Lozito, Jay R. Lieberman, Cheng-Ming Chuong, Mark S. Hurtig, Denis Evseenko

**Affiliations:** Department of Orthopaedic Surgery, Keck School of Medicine of USC, University of Southern California (USC), Los Angeles, CA, 90033, USA; Ontario Veterinary College, Department of Clinical Studies, University of Guelph, Canada; Department of Pathology, Keck School of Medicine of USC, University of Southern California (USC), Los Angeles, CA, 90033, USA; International Research Center of Wound Repair and Regeneration (iWRR), National Cheng Kung University, Tainan, Taiwan; Broad Center for Stem Cell Research and Regenerative Medicine Keck School of Medicine of USC, University of Southern California (USC), Los Angeles, CA, 90033, USA; UCLA Department of Orthopaedic Surgery David Geffen School of Medicine at UCLA, University of California, Los Angeles, CA, 90095, USA

## Abstract

Adult mammals are incapable of multi-tissue regeneration and augmentation of this potential may drastically shift current therapeutic paradigms. Here, we found that a common co-receptor of IL-6 cytokines, glycoprotein 130 (gp130), serves as a major nexus integrating various context-specific signaling inputs to either promote regenerative outcomes or aggravate disease progression. Via genetic and pharmacological experiments *in vitro* and *in vivo*, we demonstrated that a signaling tyrosine 814 (Y814) within gp130 serves as a major cellular stress sensor. Mice with constitutively inactivated Y814 (F814) exhibit regenerative, not reparative, responses after wounding in skin and anti-degenerative responses in the synovial joint. In addition, pharmacological inhibition of gp130 Y814 results in regeneration of multiple tissues in several species as well as disease modification in animal models of osteoarthritis. Our study characterizes a novel molecular mechanism that, if selectively manipulated, enhances the intrinsic regenerative capacity while preventing pathological outcomes in injury and disease.

**Summary:** Gp130 Y814 signaling module serves as a cellular stress sensor responsible for hindering tissue regeneration while triggering pathological outcomes after injury.

## Introduction

Regeneration refers to a type of healing where a compensatory new growth completely restores original tissue architecture and function following damage (*1*). Regenerative capacity of an organ is influenced by regulatory networks orchestrated by local and systemic immune responses, such as activation of macrophages, after an insult (*1, 2*). During prenatal development, mammals have an extraordinary ability to regenerate tissue; unfortunately, this capability declines with age as adult injuries are usually not regenerated, but repaired, which can result in fibrotic scarring instigating loss of original tissue integrity (*3*). Thus, it is of fundamental importance to decipher molecular mechanisms controlling degeneration and fibrosis versus regenerative responses after wounding.

In high mammals, loss of regenerative ability in the adult can be due to the inability to activate this reprogramming process, which may be in turn suppressed by the dysregulated levels of inflammatory response (*4*). Controlled signaling components of the local microenvironment, such as inflammatory reactions initiated by pro-inflammatory cytokines, contribute to successful tissue regeneration (*5*). During regeneration, inflammation often precedes the actual repair of the lesion (*5*). Previous studies across a variety of injury models suggest that a dampened inflammatory response can facilitate regeneration (*4*). IL-6 family of cytokines (IL-6, IL-11, OSM, LIF, etc.), which signal through glycoprotein 130 (gp130), are pleiotropic key players in inflammatory responses following injury that are capable of promoting both, regeneration and pathogenesis, depending on the perseverance of inflammatory signal (*6–8*). It has been reported that IL-6 cytokines including IL-6 and IL-11 play an important role during limb regeneration after injury in axolotl (*9*). The regenerative capacity of IL-6 cytokines are also seen in adult mammals in various organs and tissues (*4, 10–14*). However, aberrant and prolonged activation of IL-6 cytokines is implicated in the pathogenesis of various fibrotic pathologies (*15*) and complex chronic diseases such as osteoarthritis (OA), where progressive inflammatory and degenerative changes in articular cartilage are also accompanied by the excessive fibroplasia and collagen production in synovial tissue and subchondral bone (*16*).

The bimodal ability of IL-6/g130 signaling mechanism to simultaneously enhance regeneration and promote inflammation and fibrosis is still unclear and this gap in knowledge persists. We hypothesized that different functional outcomes downstream of gp130 may be mediated by specialized signaling module(s) of this receptor and that selective manipulation of these gp130 modules will promote regeneration in lieu of inflammation and fibrosis. In the current study, we have discovered that a signaling tyrosine 814 (Y814) residue within gp130 intracellular domain serves as a major cellular stress sensor responding to IL-6 cytokine elevation in pro-inflammatory microenvironment. Via genetic experiments *in vivo*, we have also demonstrated that inhibition of this signaling module improves regenerative outcomes in multiple tissues. Lastly, we have synthesized a small molecule modulator of gp130 receptor selectively targeting gp130 Y814 signaling outputs and have demonstrated that this drug has a major disease modifying potential in small and large animal models of OA.

## Materials and Methods

### General methods

For all experiments, biological replicates (cells from independent specimens) were employed to generate data. For experiments expected to yield large differences, standard practice of using 3-5 replicates was followed.

### Cell culture and treatments

Only early passages of chondrocytes (passage 0-2) were used for experimentation to avoid de-differentiation and loss of cartilage phenotype (*17*). Fetal tissue was provided by Dr. April Pyle (UCLA). Normal human male and female adult articular cartilage tissue (n=9, 20 to 60 years old) was obtained from the National Disease Research Interchange (NDRI). Human OA samples were obtained from NDRI and Dr. Jay Lieberman (protocol HS-16-00449 approved by the USC IRB Committee). These OA samples were dissected from the tibial and femoral articular cartilage from joint structures removed during total joint replacement. The minimally affected, based on visual evaluation, cartilage regions were dissected and used for experimentation. All patients had 3–4 on the Kellgren–Lawrence grading scale for OA. Six male (60-79 years old) and three female (55 and 70 and 81 years old) specimens were used for experimentation. All donated material was anonymous and carried no personal identifiers.

Ba/F3 and ATDC5 cells were purchased from ATCC (Manassas, VA). Mouse spleen, fat and cartilage tissue were obtained from the wild type (WT) and mutant F814 animals attained from the Jackson Laboratory or USC Transgenic Core, respectively. At termination, mouse tissues of interest were dissected, harvested and digested following previously established protocols (*17*). Legs from five-month-old Yucatan minipigs were obtained from S&S Farms (Ramona, California), and cartilage was harvested and digested following previously established protocols (*17*). Cartilage explants were made using 2 mm biopsy punch (Miltex, Inc., York, PA) and wet weight of each explant determined prior to experimentation. For cell isolation cartilage tissue was digested as described previously (*17*).

Cell culture reagents were purchased from Life Technologies, Inc. (Grand Island, New York). Growth factors LIF, OSM, IL-6, IL-11, CNTF and hyper-IL6 were purchased from Peprotech (Rocky Hill, NJ). SRC (SU6656) inhibitor was purchased from Selleckchem (Pittsburgh, PA). R805 was synthesized via a fee-for-service arrangement with Chares River, UK. Media was replenished with DMEM F12 medium containing 1% (vol/vol) fetal bovine serum and 1% Penicillin-Streptomycin (vol/vol) once treatments were added.

### Statistical analysis

Numbers of repeats for each experiment are indicated in the figure legends. Pooled data are represented as mean ± SD unless otherwise indicated. Unless otherwise indicated, statistical analysis was performed using one-way ANOVA followed by the Tukey test to compare more than 2 groups or 2-tailed Student’s *t* test to compare 2 groups. *p* values less than 0.05 were considered to be significant.

### IP Western, SDS-PAGE and Western blot analysis

For transfections of gp130 WT (InVivoGen) and gp130 mutant plasmids (Thermo Scientific), cells were plated 24h prior to transfection to produce monolayers that were 60% confluent and these were transfected by using Turbo DNAfectin 3000 (cat # G3000, Lambda Biotech) according to the manufacturer’s protocol. 72h after transfection, cells were maintained in DMEM (10% FBS, 1% PSA) and were either left untreated or were stimulated with IL-6 (10 ng/mL) and/or OSM (10 ng/mL) for 24h before cell harvest and protein extraction. For immunoprecipitation assays, cell lysates were incubated with Protein G Agarose (cat # 20398, Pierce) and anti-Flag antibody (cat # 2368, Cell Signaling) at 4°C overnight. The immune complexes were sedimented, washed and separated by SDS-PAGE (see below) and further analyzed by Western blot using p-SRC (pY416-SRC) (cat# 1246F, Novus Biologicals) or NEMO (cat # 18474-1-AP, Proteintech) antibodies and normalized to gp130 (cat #bs-1459R, Bioss) or total SRC (cat # 2123, Cell Signaling). For R805 treatment, the immune complexes were sedimented, washed and separated by SDS-PAGE and further analyzed by Western blot using OSMR (cat # ab85575, Abcam), LIFR (Santa cat # 515337, Cruz Biotechnology) or p-SRC (pY416-SRC) (Novus Biologicals, cat # 1246F). Histone H3 (cat # 9715, Cell Signaling), gp130 (cat # bs-1459R, Bioss), total SRC (cat # 2123, Cell Signaling) or Flag antibody (cat #14739, Cell Signaling) were used as loading controls.

For standard western blot, treated and non-treated cells were lysed in RIPA Lysis and Extraction Buffer (Pierce, Rockford, IL) containing protease inhibitors (Pierce) followed by sonication with a 15-second pulse at a power output of 2 using the VirSonic 100 (SP Industries Company, Warminster, PA). Protein concentrations were determined by BCA protein assay (Pierce). Proteins were resolved with SDS-PAGE utilizing 4–15% Mini-PROTEAN TGX Precast Gels and transferred to Trans-Blot Turbo Transfer Packs with a 0.2-µm pore-size nitrocellulose membrane. The SDS-PAGE running buffer, 4–15% Mini-PROTEAN TGX Precast Gels, Trans-Blot Turbo Transfer Packs with a 0.2-µm pore-size nitrocellulose membrane was purchased from Biorad (Hercules, CA). Nitrocellulose membranes were blocked in 5% nonfat milk in 0.05% (v/v) Tween 20 (PBST) (Corning, Manassas, VA). Membranes were then incubated with primary antibodies p-gp130 Y814 (Evseenko lab, AbClonal), p-AKT (cat #4060, Cell Signaling), p-ERK 1/2 (cat #9106, Cell Signaling), p-Y416-SRC (cat #1246F, Novus Biologicals), p-gp130 (cat # 1453R, Bioss USA) and p-NFκBp65 (cat #8242, Cell Signaling), c-Myc (cat #5605, Cell Signaling) and/or YAP1 (cat # 4912, Cell Signaling). Histone H3 (cat # 9515, Cell Signaling) was used as loading control. After washing in PBS containing 0.05% (v/v) Tween 20 (PBST), membranes were incubated with rabbit (cat # 31460, Thermo Scientific) or mouse IgG-HRP (cat # 31430, Thermo Scientific) secondary antibodies. After washing, membranes were developed with the Clarity Western ECL Blotting Substrate (Hercules, CA). Quantification of blots was performed using ImageJ 1.44 followed by Prism9 for analysis.

### Lizard Tail Healing Model

Captive bred and raised bearded dragon lizards (*Pogona vitticeps*) were maintained at 25.5°C with 65% humidity on a 12 hr light/12 hr dark schedule with UVB lamp (Zoo Med) treatment during daylight hours. Care and experimental use of lizards was conducted in accordance with USC Institutional Animal Care and Use Committee (IACUC) approved protocol 20992. Lizard tails were amputated halfway along lengths with scalpels To systemically deplete macrophage populations, lizards received intraperitoneal (IP) injections of L-α-phoshatidylcholine/cholesterol liposomes containing clodronate (0.125 mg/g) as previously described [Londono et al. J Immunol Regen Med. 2020]. Control animals were treated with liposomes containing PBS instead of clodronate. Lizards were treated with R805 drug via daily direct injection (1 pmol drug) into amputated tail stumps. Lizard tail samples were collected 14 days following original tail amputation. Collected tail samples were photographed using an Olympus SZX16 stereo microscope. Images were uploaded in ImageJ (NIH, Bethesda MD) and quantified using the measure command. For labeling of proliferating cell populations with 5-ethynyl-2’-deoxyuridine (EdU) (ThermoFisher Scientific), animals received IP injections of EdU (50 mg/kg) four hours prior to sample collection. Samples to be analyzed via histology were fixed overnight in 4% paraformaldehyde, decalcified for 3 weeks in 10% EDTA (pH 7.4), equilibrated to 30% sucrose, embedded in optimal cutting temperature compound (OCT, Tissue-Tek), crypsectioned at 16 µm thickness, mounted on glass slides, and stained with 4′,6-diamidino-2-phenylindole (DAPI). Samples were stained with Masson’s trichrome (American Mastertech) according to the manufacturer’s instructions or immunostained for CTSK (Abcam antibody ab19027, used at 1:1000). Images were captured with an Olympus CKX41 microscope outfitted with a Keyence BZ-X800 microscope, and the areas of EdU signal were quantified with ImageJ. Statistical analysis was performed using Prism 7 with one or two-way ANOVA with pairwise Tukey’s multiple comparison test for data with multiple groups. A p-value of <0.05 was deemed to be statistically significant. All values and graphs are shown as mean ± SD.

### RNA sequencing library preparation and sequencing

Total RNA was isolated using QIAGEN RNeasy Mini kit and quantified using Qubit fluorometer (Thermo Fisher Scientific). Quality of the isolated RNA was checked using Agilent Bioanalyzer 2100. Universal Plus mRNA-Seq Library with NuQuant (TECAN) was used to generate stranded RNA-seq libraries. Briefly, poly(A) RNA was selected followed by RNA fragmentation. Double stranded cDNA was generated thereafter using a mixture of random and oilgo(dT) priming. The library was then constructed by end repairing the cDNA to generate blunt ends, ligation of Unique Dual Index (UDI) adaptors, strand selection and PCR amplification. Different adaptors were used for multiplexing samples in one lane. Sequencing was performed on Novaseq SP with paired end 50 base pair reads. Data quality check was done on Illumina SAV. Demultiplexing was performed with Illumina CASAVA 1.8.2.

### RNA Sequencing Data Analysis

Raw fastq files were analyzed in Partek flow (version 10.0.21.0801). Reads were aligned to mouse GRCm38 (mm10) genome using Gencode Release M25 reference using STAR aligner (version 2.7.3a) (*18*). Transcript levels were quantified to the reference using Partek E/M with default parameters. Normalization was done using counts per million (CPM) method. Genes were considered to be differentially expressed based on fold change>2 and p-value<0.05. Functional enrichment analysis for the differentially expressed genes was performed using Ingenuity Pathway analysis (IPA, Qiagen). Pathway schematics were generated using Path designer application of IPA. MA-plots for the differentially expressed genes were generated in R using ggmaplot function of ggpubr (v0.4.0) package.

### Single-cell sequencing using 10X Genomics

Single cell samples were prepared using Chromium Next GEM Single Cell 3’ Reagent Kits v3.1 or Next GEM Single Cell 5’ v2 (dual index) and Chip Kits (10x Genomics) according to the manufacturer’s protocol. Briefly samples were FACS sorted using DAPI to select live cells followed by resuspension in 0.04% BSA-PBS. Nearly 1200cells/µl were added to each well of the chip with a target cell recovery estimate of 1000 cells. Thereafter Gel bead-in Emulsions (GEMs) were generated using GemCode Single-Cell Instrument. GEMs were reverse transcribed, droplets were broken and single stranded cDNA was isolated. cDNAs were cleaned up with DynaBeads and amplified. Finally, cDNAs were ligated with adapters, post-ligation products were amplified, cleaned-up with SPRIselect and purified libraries were sequenced on Novaseq at the UCLA Technology Center for Genomics and Bioinformatics.

### 10X Sequencing data analysis

Raw sequencing reads were processed using Partek Flow Analysis Software (build version 10.0.21.0210). Briefly, raw reads were checked for their quality, trimmed and reads with an average base quality score per position>30 were considered for alignment. Trimmed reads were aligned to the mouse genome (mm10) or dog genome (canFam3) using STAR-2.7.3a with default parameters. Reads with alignment percentage >75% were deduplicated based on their unique molecular identifiers (UMIs). Reads mapping to the same chromosomal location with duplicate UMIs were removed. Transcripts were quantified using mm10 Gencode vM25 (mouse) or canFam3 Ensembl v102 (dog) and UMI count matrix was generated with rows representing genes and columns representing each detected cell. Thereafter ‘Knee’ plot was constructed using the cumulative fraction of reads/UMIs for all barcodes. Barcodes below the cut-off defined by the location of the knee were assigned as true cell barcodes and quantified. Further noise filtration was done by removing cells having >5% mitochondrial counts. Genes not expressed in any cell were also removed as an additional clean-up step. Cleaned-up reads were normalized using counts per million (CPM) method followed by log-transformation generating count matrices for each sample. Count matrices were used to visualize and explore the samples in further details by generating UMAP plots, a non-linear dimensional reduction technique. This algorithm learns the underlying manifold of the data and places similar cells together in a low-dimensional space. K-means clustering was computed for identifying groups of cells with similar expression profiles using Euclidean distance metric based on the most appropriate cluster count with a maximum of 1000 iterations. Gene ontology and KEGG pathway enrichment analysis for the differentially expressed genes was performed using DAVID Gene Functional Classification Tool (http://david.abcc.ncifcrf.gov; version 6.8). Dot plots were generated in R (v4.0.3) using ggplot2(v3.3.3).

### Data availability

All RNA-seq data are deposited in GEO under the superseries accession GSE168395.

### Quantitative Real-Time PCR

Power SYBR Green (Applied Biosystems) RT-PCR amplification and detection was performed using an Applied Biosystems Step One Plus Real-Time PCR machine. The comparative Ct method for relative quantification (2-ΔΔCt) was used to quantitate gene expression. TBP (TATA-box binding protein) was used for gene normalization and expressed relative to a calibrator (sample in each set with lowest expression). Q-PCR for was conducted using the following primers: human: TBP forward, 5’ GATGGACGTTCGGTTTAGG 3’; TBP reverse, 5’ AGCAGCACAGTACGAGCAA 3’, MMP13 forward, 5’ ACTGAGAGGCTCCGAGAAATG 3’; MMP13 reverse, 5’ GAACCCCGCATCTTGGCTT 3’, ADAMTS5 forward 5’ GAACATCGACCAACTCTACTCCG 3’; ADAMTS5 reverse, 5’ CAATGCCCACCGAACCATCT 3’, ADAMTS4 forward 5’ GCAACGTCAAGGCTCCTCTT 3’; ADAMTS4 reverse, 5’ CTCCACAAATCTACTCAGTGAAGCA 3’, Collagen II forward, 5’ CCTGGCAAAGATGGTGAGACAG 3’; Collagen II reverse 5’ CCTGGTTTTCCACCTTCACCTG 3’; Aggrecan forward, 5’ AGGCAGCGTGATCCTTACC 3’; Aggrecan reverse, 5’ GGCCTCTCCAGTCTCATTCTC 3’. Cells were treated with 10 ng/mL OSM and/or with 5 µM SRC (SU6656) inhibitor, 10ng/mL OSM and/or 50ng/mL WT/F814 plasmids; 10ng/mL OSM and/or 100 or 300 ug/mL of peptide QQpYF (Evseenko lab, Thermo Scientific); 10uM R805 and/or 10ng/mL OSM.

### Cloning and transfection of Ba/F3 and ATDC5 cells

A plasmid encoding full-length gp130 (WTgp130; InVivoGen) was used to create the 5 mutants. The parental vector was digested with EcorI and NheI; 5 different DNA fragments encoding point mutations for tyrosine to phenylalanine (residue Y814), lysine to arginine (residue K816), and serine to alanine (residues S820, S824, S825) within 5’ GATGGTATTTTGCCCAGGAGTCAGCATGAATCCAGT 3’ d812-827a were purchased from Thermo Scientific and ligated into the backbone. All independent transfectants were sequenced to verify the point mutation substitution and transfected into Ba/F3 cells or ATDC5 cells. 48 hours after transfection of either WT gp130 or 5 mutants, cells were treated with or without 10 ng IL-6, OSM and/or 5uM of SRC inhibitor SU6656 and then harvested for Western blot.

### Immunohistochemistry

Histological staining for was performed on WT mice, F814 mice and all mice that underwent destabilization of the medial meniscus (DMM) (*19*). Six weeks after surgery, mice were sacrificed and joints were fixed in 10% formalin, decalcified in 10% EDTA and embedded in paraffin. For rat model, 6 weeks after partial medial meniscectomy (PMM), rats were sacrificed and joints were fixed in 10% formalin, decalcified in 10% EDTA and embedded in paraffin. A total of three 8µm thick sections were made at a 200µm interval by coronal sectioning. For canine model, 4 and 16 weeks after medial meniscal release (MMR), dogs were sacrificed and joints were fixed in 10% formalin, decalcified in 10% EDTA and embedded in paraffin. No animals were excluded from analysis. A microtome (Leica) was used to cut 5-µm sections for joints. H&E staining was performed to assess morphology. Safranin O/Fast Green staining, immunohistochemistry for proteolytic enzymes and Osteoarthritis Research Society International (OARSI) scoring was performed as described (*20, 21*). For OARSI scoring, observers performing the analysis were blinded as whether the slides were from treated or control animals. The stained samples were visually observed using LSCM (Zeiss LSM710, Carl Zeiss, Germany).

### Neoepitope analysis

WT and F814 mouse articular cartilage explants were treated with or without OSM (10 ng/mL). Pig articular cartilage explants were stimulated with or without OSM (10 ng/mL) and treated with or without peptide QQpYF (100 µg/mL or 300 µg/mL). For R805 treatment, pig articular cartilage explants were stimulated with or without OSM (10 ng/mL) and treated with or without R805 (0.01, 0.1, 1 or 10 µM). The explants were then digested for 2 hr at 37°C with 0.01 units chondroitinase ABC (Sigma Aldrich, St. Louis, MO). Samples were then dialyzed with ultrapure water for 24 hr at 4°C, freeze dried, dissolved in RIPA Lysis and Extraction Buffer (Pierce, IL) containing protease inhibitors (Pierce) followed by sonication. Proteins were separated and analyzed by Western blot using primary antibodies against aggrecan (cat# NB100-74350, Novus Biologicals,) and collagen II neoepitopes (cat #50-1035, Ibex). Wet weight of explants was used as loading control.

### Peptide QQpYF synthesis

Amino acid sequence of human gp130 was used and peptides were synthesized surrounding tyrosine 814. The peptide sequences synthesized *in vitro* in a cyclic manner were as follows: RQQYFKQNCSQHESS, DGILPRQQYFKQNCS, YFKQNCSQHESSPDIS, GDGILPRQQYFKQN, QYFKQNCSQHESSP, VDGGDGILPRQQYFK, DGGDGILPRQQYFK, QYFKQNCSQHESSPD, DGGDGILPRQQYFKQN, GDGILPRQQYFKQNC, GGDGILPRQQYFKQN, GILPRQQYFKQNCSQ, PRQQYFKQNCSQHE, PRQQYFKQNCSQHES, QQYFKQNCSQHE,QQYFK, QQYF, QYFK, YFK, QQY, and QYF. (4mg, purity >=50%) with >70% conjugation rate of peptide to protein carriers. All tyrosines (Y) were modified to phosphorylated tyrosines. The peptides were synthesized and provided by Thermo Scientific (Rockford, IL).

### Molecular docking and Molecular Dynamics Simulations

The molecular docking of Gln-Gln-phosphoTyr-Phe (QQ[pY]F) to c-Src kinase was performed in two steps. The coordinates for the c-Src Kinase was obtained from the protein data bank (PDBID 2SRC) (*22*). In the first step, protein-peptide docking server was used to identify the preferred binding site in the c-Src kinase for the QQYF peptide. For the blind c-Src Kinase-QQ[pY]F docking, HPEPDOCK server (http://huanglab.phys.hust.edu.cn/hpepdock/) was used (*23*). HPEPDOCK server uses a hierarchical flexible-peptide docking protocol, which includes conformational sampling of the peptide and peptide docking. In the second step, refinement of the protein-peptide docking was done using the GOLD, version 5.8.1 (*24*) using the best ranked site for QQ[pY]F binding to SRC kinase that was identified by the HPEPDOCK server as a starting model. All the residues within 18 Å of centroid around the initial identified site were defined as part of the peptide binding site. The GOLD refinement protocol includes 300 genetic algorithm run, 100 000 iterations were employed in which early termination option was disabled. The best docked pose was then chosen based on the GOLDSCORE and all poses were retained. Molecular dynamics (MD) simulations was carried out to determine the binding free energy of QQ[pY]F peptide binding to cSrc Kinase. AMBER18 package was used to perform the MD simulation. Amber-compatible parameters for post-translational modified amino acids were taken from Forcefiled_PTM (*25*). After adding hydrogens, the cSrc Kinase-QQ[pY]F complex was solvated in a truncated octahedral TIP3P box of 12 Å, and the system was neutralized with sodium ions. Periodic boundary conditions, Particle Mesh Ewald summation and SHAKE-enabled 2-femto seconds time steps were used. Langevin dynamics temperature control was employed with a collision rate equal to 1.0 ps^−1^. A cutoff of 13 Å was used for nonbonding interactions. Initial configurations were subjected to a 1000-step minimization with the harmonic constraints of 10 kcal.mol^-1^A°^-2^ on the protein heavy atoms. The system was gradually heated from 0 to 300 K over a period of 50 ps with harmonic constraints. The simulation at 300 K was then continued for 50 ps during which the harmonic constraints were gradually lifted. The system was then equilibrated for a period of 500 ps before the 100 ns production run. The MD simulation was carried out in the NPT ensemble. Equilibration and production run were carried out using the Sander and PMEMD modules (optimized for CUDA) of AMBER 18.0 (ff14SB) (*25*). All analyses were performed using the cpptraj module of AmberTools 18. From the 100 ns simulation 10000 structures were taken at an interval 10 ps for the free energy calculations. The MMPBSA module in AMBER was used to compute the binding free energy for QQ[yP]F binding to cSrc kinase.

### Generation of F814 CRISPR/Cas9 mouse

**Selection and synthesis of Cas9 mRNA and sgRNA (5’** AAAATGTGAAATCTCTG-GACAGG-3’) **to gp130 target region was provided by PNA Bio (**Thousand Oaks, CA) **and** targeting efficiency of the sgRNAs used for the knock-in experiment was evaluated by surveyor nuclease assay to detect the sgRNA with highest DNA cleavage efficiency. Microinjections by USC Transgenic Core were performed at one-cell stage embryo using C57BL/6 mouse strain. Mice genotyping was performed by GeneWiz (La Jolla, CA).

### Destabilization of the medial meniscus (DMM) mouse model of osteoarthritis

All experiments were conducted in accordance and under the supervision of the University of Southern California Department of Animal Resources. Six 3-month-old WT (Charles River, USA) and F814 mice (University of Southern California, USA) were anesthetized and medial para-patellar arthrotomy was carried out under a dissection microscope to expose the meniscus via knee flexion; once located, the meniscus ligament was cut, but not removed, to destabilize the joint (*19*). Only males were used in this study; females are resistant to this injury type of injury. Six weeks after the initial surgery, mice were sacrificed and the joints were transferred to 10% formalin for 48 hours followed by a decalcification process and embedding in paraffin for histological assessment.

### Wound-induced hair neogenesis assay

The WIHN assay was previously described (*26*). Briefly, a 1.5 x 1.5 cm square full thickness wound was excised on the posterior dorsum of 6-week-old mice (WT and F814), and observed for hair neogenesis and wound histology. Mice were anesthetized using isofluorane, full thickness skin was excised, and analgesic Buprenorphine SR (0.5 mg/kg) was given by intraperitoneal injection (IP) at the beginning of the procedure. Additional DietGel Boost (ClearH2O) was placed on the bottom of the cage during the first week post-operation. For R805-treatment, R805 (10 µM or 20 µM) was applied topically to the wound of WT mice.

### Alkaline phosphatase (ALP) stain

To detect newly forming dermal papillae, alkaline phosphatase staining was performed as previously reported (*26*). Briefly, full thickness wounds were excised and epidermis separated from the dermis using 20 mM EDTA. The dermis was fixed in acetone overnight at 4°C, and washed in PBS several times. The dermis was pre-incubated in ALP buffer (0.1 M Tris-HCl, 0.1 M NaCl, 5 mM MgCl_2_ and 0.1% Tween-20) for 30 min, incubated with BCIP/NBT Color Development Substrate (Promega, Madison, WI, USA) in ALP buffer at 37°C until color development. The reaction was stopped by washing with pH 8.0 Tris-EDTA and the tissue stored in PBS with sodium azide. 5 wounds were calculated for each strain of mice.

### Hair fiber length quantification

To quantify regenerated hair fiber length, hair fibers from respective wounds were plucked, aligned and photographed next to a ruler under the dissecting microscope. The length of the hair fibers was measured and analyzed using ImageJ. 3-4 hairs were randomly plucked from each wound and analyzed.

### Synthesis of phospho-gp130 Y814 antibody

Phospho-gp130-Y814 antibody was synthesized and provided by AbClonal (Woburn, MA). Synthesis of antigen modified (phosphorylated) peptide: PRQP(pY)FKQNC (19mg, purity>=85%), non-modified peptide: PRQPYFKQNC (14mg, purity>=85%), and conjugation of modified peptide to KLH was performed followed by immunization of 3 New Zealand rabbits and sera collection. Antibody purification was conducted by antigen affinity chromatography followed QC via dot-blot test against the modified and non-modified polypeptides.

### Receptor competition assay

Pig articular chondrocytes were transfected with gp130-Flag 72 hours before incubation with 0.3, 1, 3, 10 or 30 µM R805 in the presence of 10 ng/mL OSM. After 24 hours of cytokine treatment, protein was extracted and immunoprecipitated with gp130-Flag (cat # 125623, Thermo Scientific). Western blots were then performed for detection of OSMR (cat # ab85575, Abcam) and antibodies and normalized to gp130-Flag (cat # 125623, Thermo Scientific).

### Rat model of osteoarthritis

All experiments were conducted in accordance and under the supervision of the University of Southern California Department of Animal Resources. 24 male Sprague-Dawley (10 weeks old) were purchased through Charles River, USA. Animals were pair housed in standard cages in a temperature and humidity regulated room on a 12h dark/light cycle. 8 Rats received medial meniscal tear (MMT) surgery (*19*) and weekly injections of the vehicle (50 microliter of Saline + DMSO 1:1000), another 8 rats received MMT surgery and 50uL at 10 µM of R805 and 3 rats were used as sham operated control. All surgeries were performed on the right hind paw. Rats received weekly injections with either the drug or the vehicle and an empty needle in the case of the shams. Rats were anesthetized using 5% Isoflurane (VetOne inc., Boise, ID, USA) in 100% oxygen and maintained with 2% Isoflurane throughout the surgery. The joint was held in a 90-degree angle to open the joint space. Soft tissue was bluntly dissected to expose the meniscus and carefully cut to excise ∼50%. Rats were sacrificed 6 weeks’ post-surgery, joints were harvested and kept in a PBS drenched cloth in a humid chamber at 4 degrees Celsius for a maximum of 8 hours. Indentation was performed on fresh joints, and once finished, the joints were transferred to 10% formalin for 48 hours followed by a decalcification process and embedding in paraffin for histological assessment.

### Indentation and thickness mapping

Mechanical properties were mapped *ex vivo* using a 17N multi-axial load cell and a 0.5mm spherical indenter. The Mach-1 v500css, a novel developed device by Biomomentum Inc., Laval, Canada, was utilized for cartilage indentation and thickness mapping on rat articular cartilage. A camera system provided by Biomomentum was used to superimpose a position grid that was used as a template to predefine 40 positions per tibia, 40 per femur and 20 per patella. Indentation was performed at a speed of 50 µm/s using an amplitude of 50 µm, scanning grid was set on 100 µm. When indentation was finished, the spherical indenter was replaced with a 26G ½” intradermal bevel needle to determine the thickness of the predefined positions to calculate the instantaneous modulus using the model developed by Hayes (*27*). Instantaneous modulus (IM) and thickness data is presented as mean +/- SD. IM data shown represents the instantaneous modulus at 20% strain to mimic its compression *in vivo*.

### Pharmacokinetic assessment of R805

Pharmacokinetics were conducted in a canine model for over 90 days. Purpose-bred beagles with intact stifles were injected intra-articularly with 500 µL of physiological saline containing 0.25% carboxymethyl cellulose (CMC) and either 1 µg or 100 ng of CX-011 (n = 4 each). Synovial fluid and plasma were collected at d0, d3, d7, d14, d21, d31, d60 and d90 and analyzed by LC-MS/MS. Doses of 10 µg, 1 µg and 0.1 µg of R805 were utilized for the main experiment.

### Canine osteoarthritis model

All procedures were approved by an institutional animal care review board using national guidelines governing research animal welfare. Twenty-four purpose bred Foxhound cross dogs (12 females, 12 males), 10 months of age (Marshall BioResources (North Rose, NY, USA)) were chosen based on ability to walk on a leash and social interaction with handlers. Littermate information was obtained to ensure littermates were placed in separate groups to control for genetic similarities. Dogs were allocated to groups to approximate similar body weight and gender distributions. Additional daily socialization and outdoor exercise was done with trained handlers, and weekly training on an obstacle course were done for four weeks. Two weeks pre-operatively all dogs underwent baseline assessments including: lameness scoring (0 to 5) where 0 is normal and 5 is non-weight bearing, Colorado Acute Pain Scale scoring (0–4), and gait analysis using a Tekscan Walkway7 (Tekscan Inc., Boston, MA, USA). Five valid gait trials were completed for each dog at each time point. A trial was considered valid when three complete gait cycles were recorded and velocity was within 1.6-1.9 ± 0.5 m/s (Brown et al., 2013). All data was collected using Tekscan Walkway Research Software (ver. 7.66-05) and forces were normalized to each dog’s body weight (kg). Maximum force (kg) and maximum peak pressure (kPa) for each limb as well as maximum force ratios between front/hind, left front/right front and left hind/right hind were selected for analysis. At week 0 CT imaging of both knee joints was done under general anesthesia which was followed by unilateral arthroscopic transection of the caudal medial meniscotibial ligament. This medial meniscal release procedure (MMR) resulted in loss of meniscal loading sharing in order to induce OA in one knee. Dogs were allocated into four experimental groups (n=6). Three groups received CX-011 by intra-articular injection into the operated limb and the fourth control group (n=6) received an equal volume of vehicle. Intra-articular treatments were done four weeks postoperatively and repeated a second time at week 11. The three experimental groups given the test article received doses of concentration of 1 µg, 0.1 µg or 0.01 µg in a 1.0 mL volume. Knee joint CT imaging was repeated at 4 weeks and again at 16 weeks to assess OA progression. At 18 weeks the dogs were euthanized using pentobarbitol and the knee joints were harvested. MicroCT imaging at 45µ resolution using a GE eXplore Locus platform (GE Healthcare/EVS Corp., London, ON, Canada) allowed 3D reconstructions to be created and adaptive regions of interest 3 mm deep were made in the central non-meniscus covered portion of the tibial plateau subchondral bone plate. Bone morphometry parameters including tissue mineral density and trabecular bone volume were calculated using Microview™ analytical software (Parallax Innovations Ltd., London, ON, Canada). Following this the knee joints were dissected for macrophotography and biomechanical indentation testing. Thickness and instantaneous modulus were calculated for X points on the medial tibia and Y points on the medial femoral condyle using a 1 mm spherical indentor, a 0.2mm/s rate with a 5 second relaxation time. The spherical indenter was replaced with a 26 G 3/8” precision needle and an automated needle penetration test (0.5mm/s up to a predefined load to reach subchondral bone) was performed at the same pre-defined mapping points to determine cartilage thickness. Heat scale color scale maps of modulus and thickness were created. Collection of synovial tissues for histology included synovial membrane tissue sections that were stained with H&E and osteochondral sections of the medial femoral condyle and tibial plateau that were stained with both H&E and safranin-O. All sections were reviewed by one investigator blind to the treatment allocations using the canine OARSI osteoarthritis score system.

## Results

### Genetic knockout of a novel gp130 modality *in vivo* results in downregulation of pro-degenerative and pro-fibrotic signaling cascades

The molecular inflammatory processes that are activated to promote regeneration in an attempt to reestablish homeostasis after an acute injury can, when overstimulated, progressively drive degeneration of tissue and fibrosis, which is a pathogenic process where connective tissue replaces normal parenchymal tissue forming a permanent scar (*28*). Inflammation mediated by IL-6 family cytokines has been identified as a driver of chronic inflammatory processes and fibrosis (*29*). To initiate downstream signaling, gp130 requires recruitment of specific proteins on its various intracellular residues (19, 20). Engagement of gp130 by IL-6 cytokines leads to activation of various downstream signaling cascades including MAPK/ERK, PI3K/AKT, JAK/STAT3 and less renowned SRC signaling (*30, 31*), but the role of each of these modules and their interactions with gp130 are context specific and is not fully understood.

First, we wanted to assess which potential residue plays the most predominant role in tissue degradation downstream of gp130, more specifically, degradation of extracellular matrix. To test this, we stimulated pig articular chondrocytes with OSM, which we have previously shown was the most pro-inflammatory IL-6 family member (*32*), followed by treatment with numerous pharmacological inhibitors to block gp130-dependent signaling. The results demonstrated that relative to other pathways, inhibition of SRC kinase had the most significant effect of abolishing OSM-induced transcription of matrix-degrading genes *ADAMTS4* and *MMP13* (Fig. S1A). This was of an interest to us as previous studies reported SRC to be highly expressed in the synovium of patients with rheumatoid arthritis and constitutively activated in a rat model of collagen-induced arthritis (*33*). To determine which cytokine has the most substantial effect on SRC upregulation, pig articular chondrocytes were stimulated with various IL-6 cytokines and relative to all the cytokines tested, OSM displayed the strongest upregulation of phosphorylated (active) SRC (pSRC) (Fig. 1A). However, the specific mechanism of how gp130 regulates SRC activation was obscure although it is well-recognized that IL-6 ligands induce specific post-translational modifications in gp130 residues to activate signaling (*34*).

**Figure 1.**
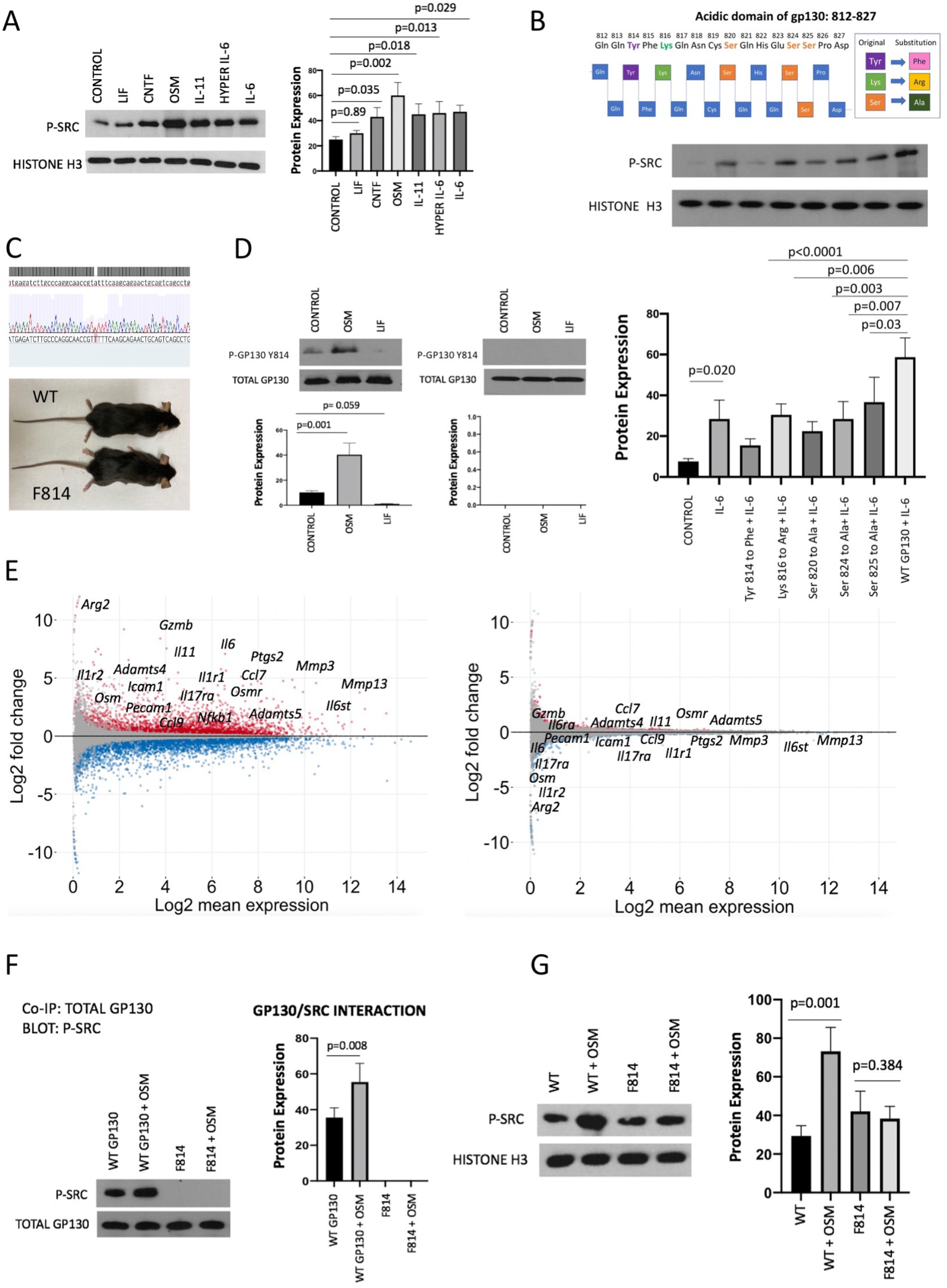
Genetic knockout of a novel gp130 modality *in vivo* results in downregulation of pro-degenerative and pro-fibrotic signaling cascades. **A.** Protein expression of pSRC in pig articular chondrocytes stimulated with IL-6 cytokines. Horizontal lines with bars show the mean ± SD. n=3. **B.** Schematic of modified amino acids within gp130 812-827 domain. Protein expression of pSRC in Ba/F3 cells after transfection with the modified or WT gp130 plasmids stimulated with IL-6 for 24 hours. Horizontal lines with bars show the mean ± SD. n=3. **C.** CRISPR gRNA was designed by PNA Bio. Y814 mutant mouse (F814) was generated by the USC transgenic mouse core. Mice genotyping was performed by GeneWiz. Representative images of 4-month-old males shown. n=4. **D.** Protein expression of gp130 pY814 in wild type (WT) and F814 mouse splenocytes treated with and without OSM and LIF for 24 hours. Horizontal lines with bars show the mean ± SD. n=3. **E.** Distribution of differentially expressed genes in MA plot of wild type (WT) and F814 mouse periarticular stromal cells. MA plots (log2fold change vs log2mean expression) for WT-OSM vs WT and Y814-OSM vs Y814 mouse periarticular stromal cells. Each dot represents a gene. Red dots represent upregulated genes i.e. log2fold-change >1 and p-value<0.05. Blue dots represent downregulated genes i.e. log2fold change < −1 and p-value 0.05. Genes that do not qualify this threshold are indicated by grey dots. **F.** Levels of protein complex formation between gp130 and pSRC in wild type and F814 mouse splenocytes stimulated with or without OSM for 24 hours. Horizontal lines with bars show the mean ± SD. n=3. **G.** Protein expression of pSRC in wild type and F814 mouse splenocytes stimulated with or without OSM for 24 hours. Horizontal lines with bars show the mean ± SD. n=3.

Based on previously published data identifying acidic domain 812-827 of gp130 as the primary region of SRC activation (*31*), we were intrigued to identify the specific residue responsible for SRC signaling. In order to determine which specific residue was responsible for this activation within residues 812-827, we have modified various amino acids within this domain through sequence substitutions using plasmids *in vitro*. Using Ba/F3 cell line, which is completely deficient of gp130, we have transfected the cells with plasmids containing wild type (WT) gp130 or gp130 sequence substitutions and stimulated with IL-6 to induce gp130 signaling. The results delineated that substitution of tyrosine (Y) to phenylalanine (F) at residue Y814 (F814) significantly reduced IL-6-stimulated pSRC (phosphorylated) expression relative to the control WT gp130 plasmid suggesting that Y814 might be responsible for pSRC regulation (Fig. 1B). Subsequently, we wanted to verify that mutation in Y814 can prevent endogenous recruitment of SRC to gp130 blocking activation of matrix degrading enzymes. To demonstrate this, we used an ATDC5 cell line, which is a chondrogenic cell line where gp130 expression is minimal. Transfection of ATDC5 with F814 plasmid stimulated with OSM markedly reduced pSRC stimulation relative to the control WT gp130 plasmid (Fig. S1B), and in the same cells, there was a decreased expression of *MMP13* and *ADAMTS4*, and an increased biosynthesis as confirmed by an increase in collagen type II (*COL2*) transcription (Fig. S1C). As such, tyrosine to phenylalanine modification of Y814 confirmed the necessity of this residue for SRC activation in chondrocytes suggesting that Y814 may play a role in cytokine-induced pro-inflammatory and pro-fibrotic signaling cascades downstream of gp130.

To employ as a tool for detection of Y814, we have developed a polyclonal antibody against phosphorylated (active) gp130 Y814 (pY814) (see Materials and Methods). Since we have previously shown that Y814 may potentially induce biosynthesis, we wanted to see if there was a difference in expression of Y814 in a developing anabolic fetal joint verses adult joint. As expected, pY814 activation was significantly lower in fetal chondrocytes compared to adult (Fig. S1D) suggesting that Y814 does not play a major role in rapidly proliferating and highly anabolic primary cartilage cells.

In order to expose the function of this residue *in vivo*, we have generated, for the first time, a CRISPR/Cas9 homozygous murine model with a genetically modified gp130 Y814 (F814). The mouse was validated by functional tests and genetic sequencing (see Materials and Methods). The mutant mouse is viable, fertile, and exhibits no significant morphological differences in the musculoskeletal tissues or other organs relative to the WT (Fig. 1C, Fig. S2). To validate the deletion of Y814 in the mutant mice, we utilized F814 mouse splenocytes and confirmed that no Y814 was detected endogenously or in response to OSM treatment though marked activation of this residue was seen in WT cells (Fig. 1D, Fig. S3). Consecutively, we wanted to identify the potential differences in reactivity to IL-6 cytokines between the WT and F814 mouse. WT and F814 mouse periarticular fibroblast cells were treated with or without OSM and subjected to RNA sequencing (RNA seq). Ordinarily, OSM upregulates a variety of pro-inflammatory and pro-fibrotic genes. Strikingly, RNA seq results demonstrated that deletion of Y814 dramatically reduces expression of these genes, including major proteases (e.g. *Mmps 3, 13; Adamts 4,5*; *Gzmb*, cytokines and their receptors (e.g. *Il-11*, *Il-6*, *Osm, Il-1r1&2*, *Osmr*, *Il-17ra*), and major pro-inflammatory regulators (e.g. *NF-kb1*, *Ptgs2/Cox2*) in response to OSM. Ingenuity Pathway Analysis (IPA) suggested the *Role of Macrophages, Fibroblasts and Endothelial Cells in Rheumatoid Arthritis* (-log_10_p-value= 7.3 or p-value= 5.01187E-08) (Fig. S4) and other pro-inflammatory and fibrosis-related pathways being the most affected by this mutation. MA plots for differential gene expression analysis confirmed that upon treatment with OSM, WT mouse fibroblasts upregulate a variety of pro-inflammatory and pro-fibrotic genes while F814 cells do not show much change in downstream gene expression (Fig. 1E) alluding to the fact that Y814 is a major stress sensor. Sequentially, mutant-derived splenocytes revealed no endogenous interaction of SRC with gp130 as shown by co-immunoprecipitation (Fig. 1F) and no activation of pSRC after stimulation with OSM stimulation was detected (Fig. 1G).

STAT3 and YAP signaling pathways downstream of gp130 play a major role in regeneration, development, and diseases such as osteoarthritis (*35–37*). Stimulation of WT and F814 mouse primary periarticular fibroblasts with LIF suggested that gp130 Y814 residue is minimally responsible for activation of STAT3 or YAP signaling as the mutant cells were not resistant to LIF-stimulated phosphorylated (active) STAT3 (pSTAT3) or YAP1 mirroring WT cells (Fig. S5). STAT3 and YAP1 levels were evaluated in response to LIF downstream of gp130 since we have previously shown that LIF primarily signals through STAT3 in chondrocytes (*32*). It is also noteworthy to mention that in WT mouse splenocytes, treatment with LIF did not upregulate gp130 pY814 further confirming that LIF is not involved in Y814 signaling (Fig. 1D, Fig. S3). Even though groups have reported that Y814 is one of the four residues responsible for STAT3 recruitment (*38*), it may not be the primary docking site, which would explain why we observe this lack of STAT3 regulation in the F814 cells. Simultaneously, genetic ablation of Y814 has completely blunted OSM-mediated activation of SRC and MAPK/ERK signaling further supporting its role exclusively in pro-inflammatory and pro-fibrotic signaling (Fig. S5).

### Gp130 Y814-deficient mouse demonstrates increased resistance to degenerative joint disease and enhanced skin regeneration after injury

To assess the direct effect of the Y814 mutation on matrix loss, we performed a neoepitope assay (*32*) using isolated femoral head explants cultured in the presence or absence of OSM. The results confirmed a significant decrease in cartilage degeneration in femoral head of Y814 mice relative to the WT as shown by the low aggrecanase and collagenase activity in the mutant (Fig. S6). To address whether Y814 mutation can ameliorate cartilage degeneration in joint disease, we adopted a destabilization of the medial meniscus murine model (*39*). This experiment clearly showed that F814 mice are resistant to surgically-induced OA as reflected by reduced loss of proteoglycans, reduced synovitis and synovial fibrosis (Fig. 2A, Fig. S7).

**Figure 2.**
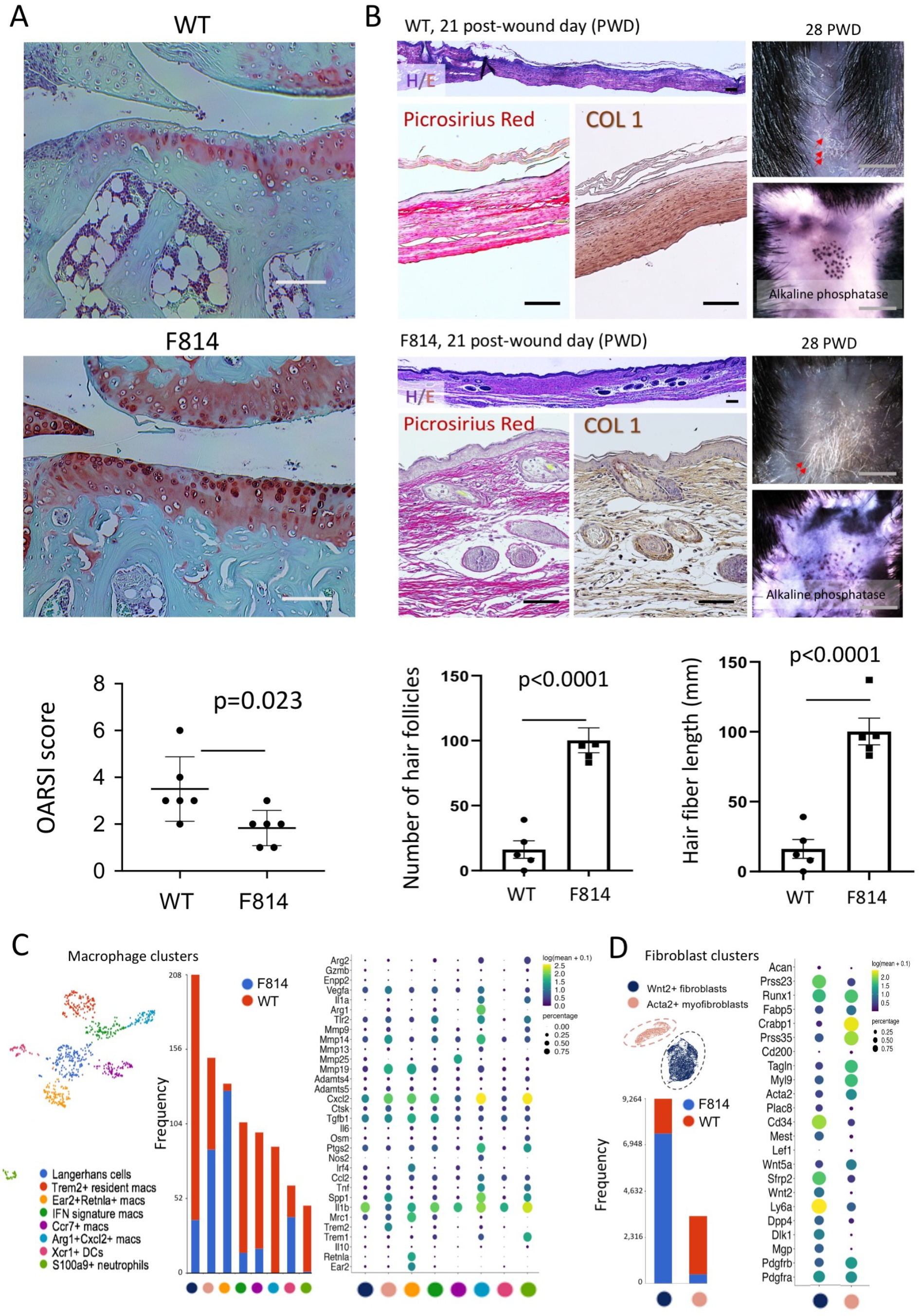
Gp130 Y814-deficient mouse demonstrates increased resistance to degenerative joint disease and enhanced skin regeneration after injury. **A.** Histological staining and quantitative assessment of cartilage degradation and changes in joint morphology of wild type (WT) and F814 mouse knee joints 6 weeks after destabilization of the medial meniscus surgery. Safranin O delineates proteoglycans (pink). OARSI scoring system was used to quantify the extent of cartilage damage. Scale bars represent 100 µm. Horizontal lines with bars show the mean ± SD. n=6. **B.** Wound-induced hair neogenesis in F814 mice is enhanced. WT and F814 mouse post-wound day (PWD) 21 wound sections after wound excision. Representative images are shown. Horizontal lines with bars show the mean ± SD. n=5. **C.** Re-clustering of clusters annotated as macrophages from WT and F814 skin wounds PWD 14. Dot plots depict gene expression in each macrophage cluster. Clusters are named based on highly significant biomarkers and gene expression profiles. The contribution of each sample to each cluster is shown as a stacked bar graph. Dot sizes are proportional to the percentage of cells in each cluster expressing the indicated gene. DCs = dendritic cells. **D.** Re-clustering of clusters annotated as fibroblasts from WT and F814 skin wounds post-wound day (PWD) 14. Dot plots depicting gene expression in each fibroblast cluster. The contribution of each sample to each cluster is shown as a stacked bar graph.

As previously mentioned, pro-inflammatory reactions resulting from tissue damage, if modulated, can contribute to successful regeneration of tissue. Since we have previously shown that ablation of Y814 prevents aberrant inflammatory signaling initiated by pro-inflammatory cytokines while bypassing the pro-regenerative signaling *in vitro*, we wanted to determine whether F814 mouse has superior regenerative potential *in vivo*. Unfortunately, OA animal models are not well-suited to study regeneration; thus, we instead employed a canonical skin excisional wound model where we accessed and compared the regenerative ability of the WT and F814 mouse skin using the wound-induced hair neogenesis (WIHN) assay (*26*) as evidence of hair follicle neogenesis and dermal regeneration. The histological observations after the procedure show, at the same magnification, that the post-wound day (PWD) 21 WT wounds are thinner and very few hair placodes are observed (Fig. 2B). Early hair placode marker P-cadherin and hair follicle bulge cells surface marker CD34 were stained negative in the wound (Fig. S8). On the contrary, the WT wound expressed high levels of notorious pro-fibrotic markers, α-SMA and COL1, and positively stained for Picrosirius red in the dermis showing high collagen deposition in addition to being contracted indicating fibrosis (Fig. 2B). In the PWD 21 F814 mouse sections, the wound is thicker and many prominent newly formed hair follicles were observed; α-SMA, CD34 and P-cadherin were expressed mostly within the hair placodes and follicles (Fig. 2B, Fig. S8). In addition, the intensity of Picrosirius red and COL1 staining in the wound dermis was lower in F814 mouse relative to the WT mouse, and the wound was not as contracted signifying less fibrosis (Fig. 2B). On PWD 28, the F814 mice showed significantly more regenerated hair follicles and longer hair fiber length than WT wounds (Fig. 2B), which reflects that the mutant wounds not only healed with a better outcome but also healed faster. Nevertheless, it is still uncertain which cell types are affected by this mutation. However, our preliminary data shows that in the periarticular subcutaneous adipose fibroblasts, F814 mouse cells are resistant to OSM-induced collagen (COL1/3) deposition unlike the WT as demonstrated by Picrosirius Red staining, which supports our *in vivo* data (Fig. S9).

To interrogate the cellular and molecular basis for the increased WIHN observed in F814 wounds, we conducted single-cell RNA sequencing (scRNA-seq) on PWD14 wounds, which is an active phase for wound healing. Live, Ter119-cells (Ter119 marks erythroid lineage) from 2-3 wounds from each genotype were sorted and processed using 10X genomics technology; cell numbers were downsampled to normalize the number of cells analyzed. UMAP followed by k-means clustering of the data delineated 9 different clusters, 4 of which expressed fibroblast genes and 2 of which were comprised of hematopoietic cells (Fig S10). Three of the fibroblast clusters and both hematopoietic clusters were disproportionately comprised of cells from one genotype (Fig S10). As macrophages have been implicated in wound healing and regenerative responses, we re-clustered the two hematopoietic clusters following exclusion of CD3+ T cells and conducted k-means clustering (Fig. 2C). This analysis yielded eight clusters which were annotated based on their biomarkers (Fig. 2C). Of note, several clusters enriched for pro-inflammatory cytokines and proteins were disproportionately populated by cells derived from WT wounds. In contrast, a cluster evidencing high levels of Ear2, Retnla and Tgfb1, genes associated with anti-inflammatory, pro-regenerative macrophages (*40*), was constituted almost exclusively by cells isolated from F814 wounds (Fig. 2C). These data indicate that in addition to fewer macrophages being present in F814 wounds, they are biased toward a less inflammatory phenotype. Re-clustering and k-means clustering of the fibroblasts from both wound types yielded two clusters, with each populated primarily by one genotype (Fig. 2D). In the cluster dominated by cells from F814 wounds, cells expressed genes associated with both papillary (Dpp4/CD26) and reticular (Dlk1) fibroblasts (*41*). Notably, this cluster was enriched for genes related to Wnt signaling including Wnt2, Sfrp2 and Sfrp4 (Fig. 2D); Wnt signaling is a known requirement for regenerative fibroblast competency in WIHN (*42*). In contrast, the cluster comprised mostly of cells from WT wounds was enriched for genes associated with myofibroblasts including Acta2, Myl9 and Tagln (*43*) (Fig. 2D). These data suggest that fibroblasts present in F814 wounds have increased competency to support regenerative vs. fibrotic wound resolution. Altogether, these results suggest that the enhanced healing observed in F814 mouse may be a result of macrophage polarization towards an anti-inflammatory phenotype and improvement of the milieu in the wound.

### An SRC-targeting peptide QQpYF prevents physical interaction of gp130 and SRC and ameliorates degenerative outcomes

Our studies delineate that gp130 Y814 regulates SRC downstream signaling; since SRC is known to associate with gp130, we hypothesized that gp130 and SRC are capable of physically interacting on Y814 and hindering this interaction will prevent SRC from signaling. In order to impede this interaction, we performed an innovative peptide library screen with short and overlapping peptides mimicking the motif of gp130 around Y814 that were individually tested for pSRC downregulation; these peptides ranged from 4-15 amino acids. The peptides contained a protein sequence stretch of consecutive amino acids surrounding gp130 Y814 residue. To test their efficacy, pig articular chondrocytes were stimulated with or without OSM in presence or absence of the peptides followed by co-immunoprecipitation to determine interaction and western blot detecting pSRC activity. Out of all the synthesized peptides, a 4-amino acid interfering peptide (QQ[pY]F), proved to be the most efficacious at physically hindering gp130-SRC protein-protein interaction in pig articular chondrocytes by binding to SRC (c-SRC), preventing accessibility to Y814 as an active site (Fig. 3A, Fig. S11A) and decreasing pSRC activation (Fig. 3B, Fig. S11B). Due to sequence specificity, the peptide is capable of discriminating among other gp130 regions serving as a competitive inhibitor. Docking for peptide–protein interaction interfaces confirmed that peptide QQpYF has a strong binding affinity to SRC as shown by the spatial proximity through protein folding in resolved 3D structure (Fig. 3C).

**Figure 3.**
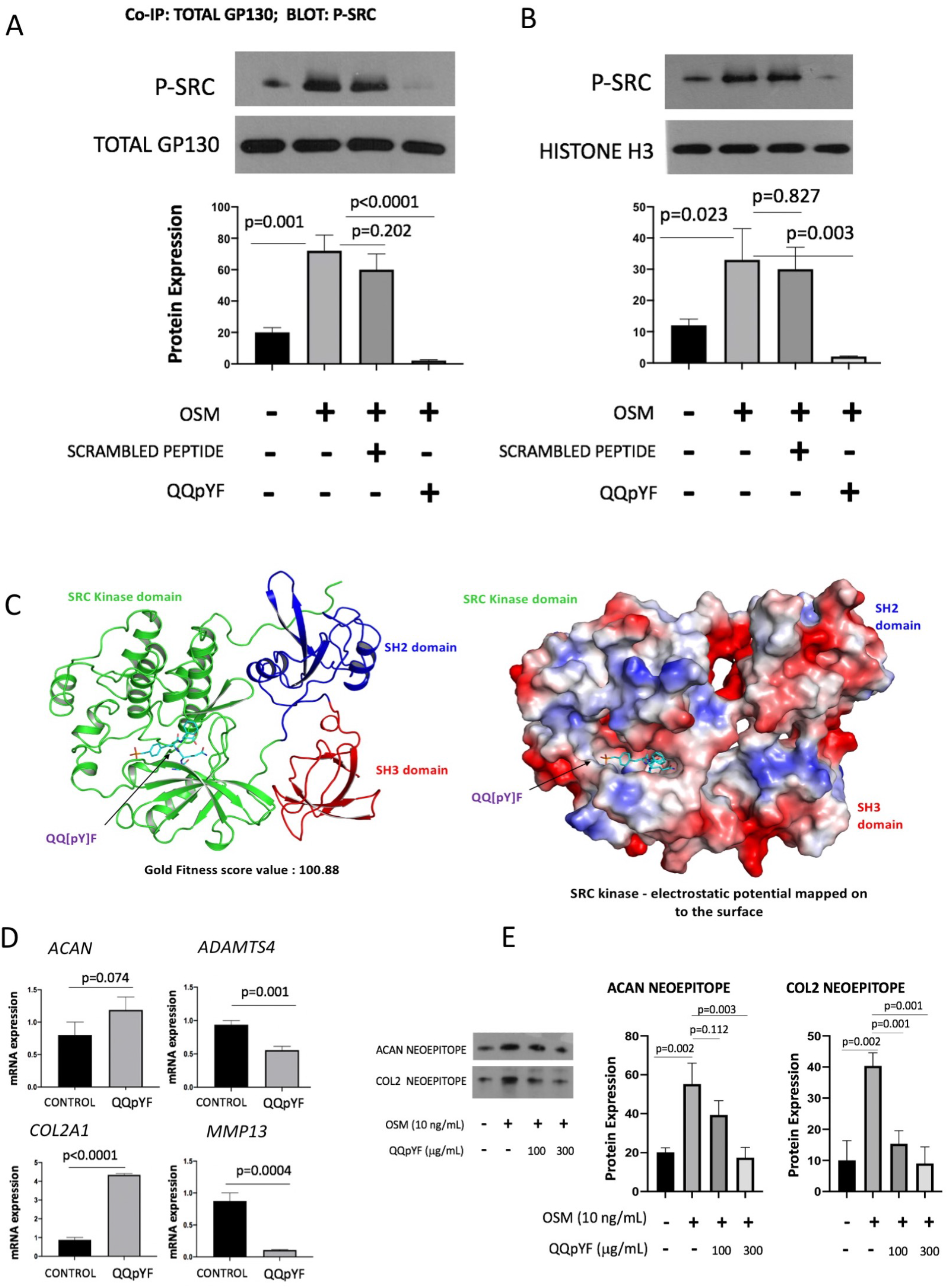
An SRC-targeting peptide QQpYF prevents physical interaction of gp130 and SRC and ameliorates degenerative outcomes. **A.** Levels of complex formation between gp130 and pSRC in pig articular chondrocytes stimulated with or without OSM in presence or absence of control scrambled peptide or peptide QQpYF for 24 hours. Horizontal lines with bars show the mean ± SD. n=3. **B.** Protein levels of pSRC in pig articular chondrocytes stimulated with or without OSM in presence or absence of control scrambled peptide or peptide QQpYF for 24 hours. Horizontal lines with bars show the mean ± SD. n=3. **C.** Computational analysis using GOLD software predicted a high affinity binding of peptide QQpYF to the regulatory site of SRC (c-SRC). c-SRC as visualized by crystallography. The structure of the indicated c-SRC domains is shown in ribbon diagram representation (left) as well as with electrostatic potential (blue, positive charge; red, negative charge; white, neutral) mapped onto the molecular surface (right). Peptide QQpYF is shown in stick representation. **D.** Transcription of genes was determined via qPCR in human adult OA articular chondrocytes treated with or without peptide QQpYF for 48 hours. Horizontal lines with bars show the mean ± SD. n=3. **E.** Pig knee cartilage explants were stimulated with or without OSM and treated with or without peptide QQpYF at indicated doses for 72 hours followed by a neoepitope assay. Levels of cleaved ACAN and COL2 neoepitopes in the supernatant were quantified with respect to the wet weight of the explant. Horizontal lines with bars show the mean ± SD. n=3.

To determine the effect of peptide QQpYF on matrix degeneration, we performed qPCR to quantify gene expression of matrix degrading enzymes in human adult OA articular chondrocytes. Transcription of *ADAMTS4/5* and *MMP13*, were markedly lower in cells treated with peptide QQpYF while biosynthesis of COL2 and aggrecan (ACAN) was increased (Fig. 3D). Further, we performed a neoepitope explant assay where pig articular cartilage explants were stimulated with or without OSM and treated with or without peptide QQpYF at different doses and the levels of cleaved ACAN and COL2 neoepitope were measured. The results confirmed the protective effect of peptide QQpYF on matrix degeneration as shown by the decrease of aggrecanase and collagenase activity (Fig. 3E).

### Gp130 Y814-targeting small molecule promotes regeneration of tissues after injury

Our previous studies in the areas of cartilage tissue degeneration and repair have identified a small molecule, RCGD 423, capable of modulating gp130 receptor signaling (*32*). RCGD 423 prevents activation of MAPK/ERK and NF-κB pathways and demonstrates strong anti-inflammatory and anti-degenerative outcomes; this molecule was also shown to highly activate STAT3 signaling and its downstream target, proto-oncogene MYC (*32*), deeming activation of this signaling potentially detrimental if hyperactivated, which is unsuitable for therapy. Therefore, it was imperative to find an analog that induces the equivalent beneficial functional outcomes but that does not upregulate MYC signaling while preventing upregulation of pro-inflammatory pathways. Importantly, our detailed analysis of human primary OA chondrocytes has demonstrated high levels of pSTAT3 (phosphorylated) upregulation compared to adult articular chondrocytes from healthy joints (Fig. S12). We have previously shown that STAT3 signaling is highly upregulated in rapidly growing, anabolic fetal chondrocytes (*32*). Thus, STAT3 upregulation in OA cartilage may be due to an intrinsic attempt for the OA chondrocytes to harness their regenerative potential making additional activation of STAT3 during disease progression, at least in context of the joint, unnecessary. Out of a large library of RCGD 423 analogs that did not upregulate MYC in response to IL-6 family of cytokines (Fig. S13), we selected an analog that we termed R805 (Fig. 4A) for further characterization as it also minimally affected STAT3 signaling in a diseased joint (Fig. S14). R805 not only markedly suppressed IL-6 family-induced heterodimerization of gp130 with their α-receptors (Fig. S15, S16) similar to RCGD 423 (*32*) and pro-inflammatory signaling in a diseased joint (Fig. S14), but it also selectively inhibited OSM-stimulated activation of Y814 (Fig. 4A, Fig. S17A). To measure the protective effect of R805 on matrix destruction, we treated pig articular chondrocytes with the drug after stimulation with OSM; based on the data, R805 was able to decrease the *MMP13* and *ADAMTS4/5* gene expression (Fig S17B) along with the levels of COL2 and ACAN neoepitopes in OSM-treated explants (Fig. S17C).

**Figure 4.**
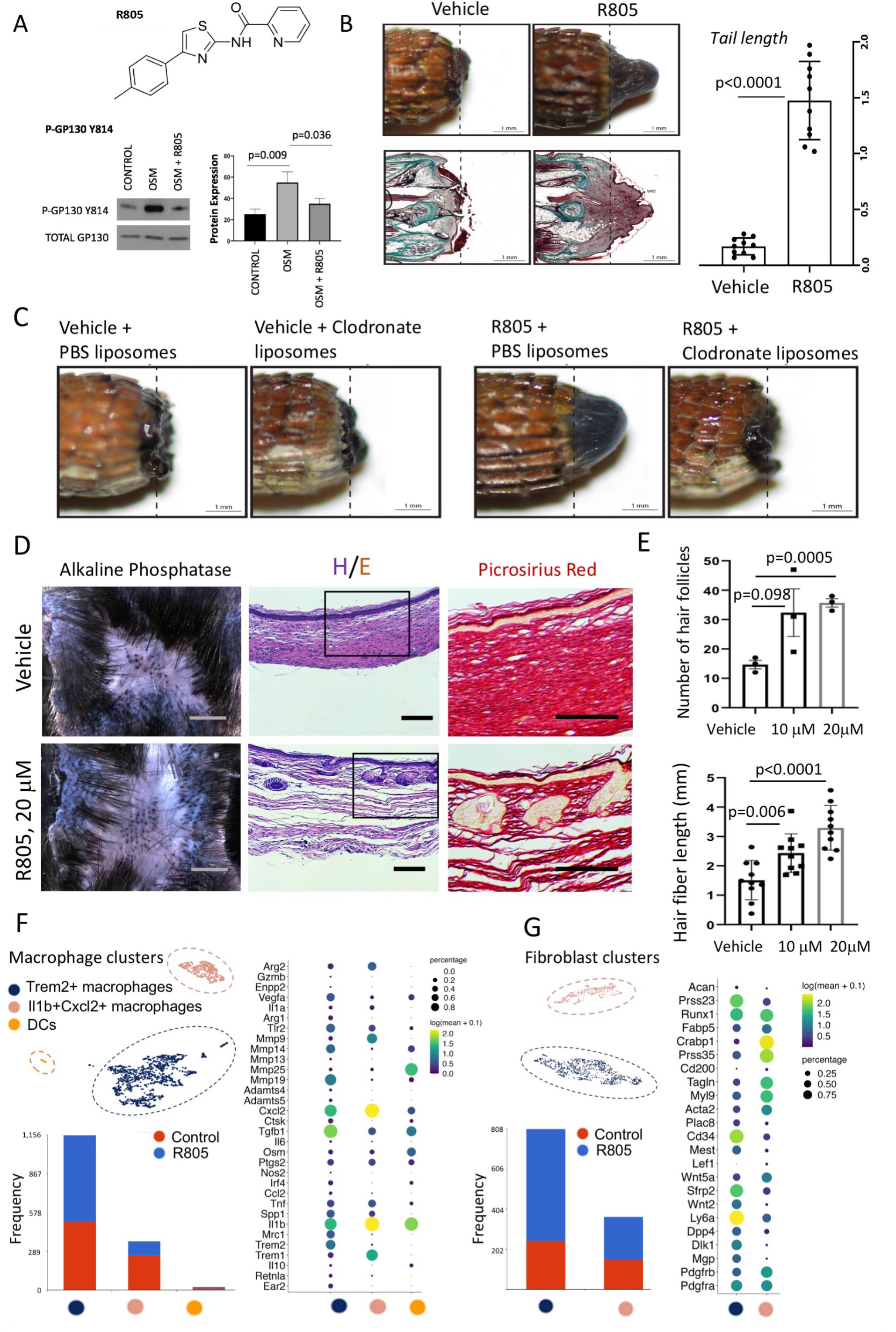
Gp130 Y814-targeting small molecule promotes regeneration of tissues after injury. **A.** Protein expression of gp130 pY814 in pig articular chondrocytes treated with or without OSM and R805 for 24 hours. Horizontal lines with bars show the mean ± SD. n=3. **B.** Gross morphology and Masson’s trichrome staining of bearded dragon treated with vehicle control and R805 14 days post-amputation. Dashed lines mark amputation planes. n=10. sc=spinal cord; we=wound epithelium; ve=vertebra. **C.** Gross morphology of bearded dragon co-treated with vehicle control or R805 and PBS or clodronate liposomes 14 days post-amputation. Dashed lines mark amputation planes. n=10. **D.** Mouse vehicle-treated or R805-treated post-wound day (PWD) 21 wound sections. Representative images are shown. n=8. **E.** Hair follicle (n=3) and fiber length (n=8) in R805-treated mouse PWD 14 and untreated control wounds. Horizontal lines with bars show the mean ± SD. **F.** Re-clustering of clusters annotated as macrophages from skin wounds of vehicle-treated or R805-treated mice PWD 14. Dot plots depict gene expression in each macrophage cluster. The contribution of each sample to each cluster is shown as a stacked bar graph. Dot sizes are proportional to the percentage of cells in each cluster expressing the indicated gene. DCs = dendritic cells. **G.** Re-clustering of clusters annotated as fibroblasts from skin wounds of vehicle-treated or R805-treated mice PWD 14. Dot plots depict gene expression in each fibroblast cluster. The contribution of each sample to each cluster is shown as a stacked bar graph.

Moreover, we wanted to determine whether R805 is inclined to shift the signaling outputs downstream of gp130 promoting differential regulation similar to genetic ablation of Y814. As a readout, protein levels of phosphorylated SRC, MAPK (ERK 1/2) and NF-κB were measured in response to OSM while STAT3 and YAP1 levels were evaluated in response to LIF. While pSRC, MAPK (pERK 1/2) and pNF-κB (pNF-κBp65) activation induced by OSM stimulation was markedly suppressed by R805 (Fig. S18), the drug had minimal inhibitory effect on pSTAT3 and YAP1 downstream of LIF (Fig. S19) in pig articular chondrocytes solidifying the role of R805 as a modulator of gp130 receptor signaling rather than a complete inhibitor. Furthermore, R805 was able to prevent endogenous interaction of SRC with gp130 (Fig. S20). These data support the concept that signaling downstream of gp130 can be parsed and modulated such that pro-degenerative activation in a pathological signaling milieu can be greatly reduced while maintaining pro-regenerative signaling supporting the data obtained from WT and F814 mouse cells.

We further wanted to test whether R805 can induce tissue regeneration in multiple animal models. First, we employed a lizard tail injury model in a lizard species not naturally capable of tail regeneration, the bearded dragon (*Pogona vitticeps*). Injection of amputated lizard tails with R805 resulted in tissue regeneration as evidenced by blastema-like structures (Fig. 4B). However, control untreated tails instead formed fibrotic scars with no new tissue besides blood clot accumulation distal to amputation sites and degenerated tissue (Fig. 4B). Masson’s trichrome staining showed that unlike R805-treated lizard tails, control tails exhibited open tail wounds at 14 days post-amputation and tail spinal cords formed pericyte scars within vertebrae centers (Fig. 4B). Since we have previously shown that lizard tail regeneration is dependent upon the recruitment of CTSK+ macrophage populations to tail amputation sites (*44*), control and R805-treated tails were analyzed by CTSK and EdU staining (Fig. S21A). Unlike control, R805-treated lizard tails exhibited CTSK+ macrophage populations that extended into new tissue areas distal to amputation planes and demonstrated significantly higher numbers of proliferating EdU+ wound epithelium and periosteal cells (Fig. S21A). These results suggested that R805 treatment increases macrophage recruitment to lizard tail amputation injuries leading to cell proliferation and tissue regeneration over fibrosis. To further solidify the role of macrophages in inducing lizard tail blastema formation and regeneration, the amputated lizard tails were co-treated with R805 with and without clodronate liposomes, which systemically deplete phagocytic macrophage populations. While lizard tails co-treated with R805 and PBS liposomes exhibited blastema-like structures, co-treatment with clodronate liposomes inhibited R805-induced tissue regrowth (Fig. 4C). Subsequently, CTSK and EdU staining showed that co-treatment with R805 and PBS liposomes exhibited recruitment of CTSK+ macrophages and EdU+ proliferative cells distal to amputation planes while treatment with clodronate liposomes inhibited this macrophage recruitment and cell proliferation (Fig. S21B). These results suggested that macrophage populations are required for R805-induced lizard tail tissue regeneration.

Since the data obtained from lizard tails was encouraging, we wanted to confirm the regeneration potential of R805 in a mammalian model. For this, we again employed a skin excisional wound model where we accessed and compared the regenerative ability of the R805-treated and untreated mouse WT skin using the WIHN assay. On PWD 21, the results demonstrated a significant hair follicle neogenesis and dermal regeneration upon topical administration of R805 on the mouse wound (Fig. 4D). The histological observations show that the untreated mouse wound is thinner; furthermore, it positively stained for Picrosirius red, α-SMA and COL1 in the dermis confirming high collagen deposition and is contracted, which indicates fibrosis (Fig. 4D, Fig. S22). This is contrary to R805-treated wound as it is thicker, less contracted and shows less collagen deposition (Fig. 4D, Fig. S22) comparable to the F814 mouse after skin excision. The increase of hair fiber length and number of hair follicles (Fig. 4E) also reveals that the treated wounds healed faster and with a better regenerative outcome, which was also concentration dependent.

Unsupervised k-means clustering defined a clear macrophage subset (Fig. S23, **Table S1**), and re-clustering of these cells delineated two macrophage clusters and a small dendritic cell cluster (Fig. 4F). The larger macrophage cluster contained more cells contributed by R805-treated wounds and evidenced higher expression of genes characteristic of pro-regenerative/anti-inflammatory macrophages including Mrc1, Trem2 and Tgfb1 (*45*) (Fig. 4F). In contrast, the smaller cluster evidenced more pro-inflammatory gene expression (Il1b, Cxcl2 and Trem1 (*46–48*)) and was populated primarily by cells from control wounds (Fig. 4F). Re-clustering of fibroblasts in R805-treated and control wounds (Fig. 4G) yielded strikingly similar results to those observed with genetic manipulation of Y814, indicating that pharmacologic modulation of this residue can support tissue regeneration by shifting the microenvironment toward a less pro-inflammatory milieu in the damaged tissue.

### R805 demonstrates disease modifying effects in small and large animal models of osteoarthritis

We then tested the ability of R805 to improve structural and functional outcomes *in vivo* in two commonly used models of posttraumatic OA. First, we adopted a rat medial meniscal tear model that has shown to be an useful for the assessment of anti-arthritic drugs *in vivo* (*49*). This study showed prominent protective effects of R805 in the joint (Fig. S24A, B). Since our *in vivo* results in rats demonstrated efficacy of R805 in alleviating cartilage degradation after injury, a preliminary 90-day pharmacokinetic study was conducted in Beagles that showed a 1 µg intra-articular (IA) injection maintained IC50 concentrations of R805 in the joint for 3 weeks; the compound was never detected in systemic circulation (Fig. S25A). Based on this data, doses of 10 µg, 1 µg and 0.1 µg IA were used in the main experiment. We conducted a high-value translational canine OA model employing baseline neuromuscular conditioning and performance assessment on stairs and obstacles (*50*), Colorado Pain Scale scoring (*51*), gait analysis and CT imaging before conducting a minimally invasive arthroscopic meniscal release (*52*) to evaluate efficacy of R805. The four experimental groups received 1 µg, 0.1 µg or 0.01 µg of R805 or saline on postoperative week 4 and 11 (Fig. S25B).

Histological abnormalities due to loss of meniscal load sharing in saline and 0.1 ug dose groups included focal partial- and full thickness cartilage erosions in central non-meniscus covered portion of the tibial plateau and longer linear erosions in the femoral condyle. In adjacent cartilage, there was a loss of the superficial collagen layer and its chondrocytes, loss of proteoglycan staining, invasion of the calcified cartilage by chondroclasts and vasculature, and some thickening (sclerosis) of the subchondral plate. There was incremental improvement in the histological scores as the R805 dose increased; erosions were more shallow and smaller, adjacent cartilage organization was preserved as was proteoglycan staining (Fig. 5A). Synovial membrane in the high and middle dose groups had mild intimal hypertrophy with occasional subintimal mononuclear cell infiltration, whereas low dose and saline treated dogs had a profound increase in subintimal vasculature, new connective tissue and cellular infiltrate comprised of lymphocytes and macrophages (Fig. 5B).

**Figure 5.**
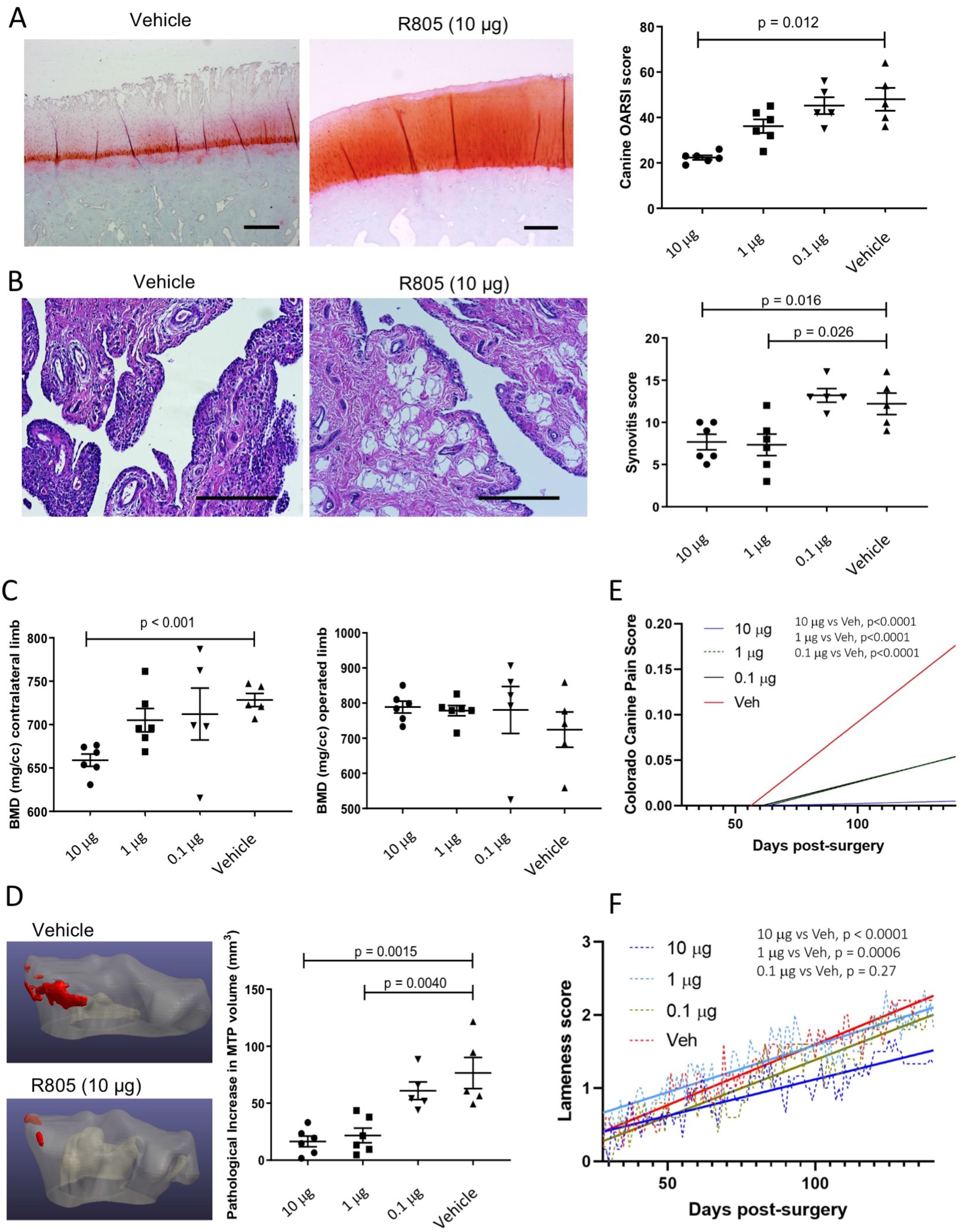
R805 demonstrates disease modifying effects in small and large animal models of osteoarthritis. **A.** OARSI scoring of canine joint sections. Horizontal lines with bars show the mean ± SEM. n=5-6. **B.** Synovial inflammation, as gauged by fibrillations and immune infiltration, is reduced by mid and high doses of R805 in canine joint. Representative images are shown. Horizontal lines with bars show the mean ± SEM. n=5-6. **C.** Assessment of canine bone mineral density (BMD) in the medial compartment of the operated stifle by microCT revealed a consistent absence of sclerosis in R805 10 and 1 µg groups. In contrast, BMD in the contralateral, non-operated medial compartment increased with decreasing dose of R805. Horizontal lines with bars show the mean ± SEM. n=5-6. **D.** Canine knee joint shape change from baseline as determined by serial CT imaging showed dose-dependent reduction in ectopic bone formation (red). Horizontal lines with bars show the mean ± SEM. n=5-6. **E.** Canine Colorado Pain Scores were measured daily and totaled for each group starting at after the first intra-articular injection. Linear regression analysis of daily scores is presented; p value. n=5-6. **F.** Linear regression analysis of lameness scores of R805-treated (10 ug) and vehicle-treated canine group. n=5-6.

Since these dogs were trained on an agility course and were leash exercised daily outdoors weather permitting, they did not develop hindleg peak vertical force asymmetry as expected. The proportion of forelimb to hindlimb weight bearing in quadrupeds is 60/40. Typically, OA drives bone remodeling and drift in shape in the medial compartment, enlarging the weight bearing surfaces, but almost no change in the cortical outline occurred in the medial tibial plateau in the dogs receiving the 10 ug and 1 ug doses. As expected, bone mineral density increased in the medial tibial plateau in response to meniscal deficiency, but more intriguingly, dogs receiving high dose R805 did not develop subchondral sclerosis in the contralateral knee that usually arises from compensatory load transfer (Fig. 5C). Knee joint shape change, a validated imaging biomarker of OA in people (*53*) and animals (*54*) was also clearly controlled in dogs receiving IA R805 (Fig. 5D). This lack of the joint shape in the high dose group was consistent with pain scoring and gait analysis data that show the high dose group had minimal pain (Fig. 5E) and lameness (Fig. 5F).

Finally, to further confirm the histological findings of reduced cartilage degeneration and synovial fibrosis, we conducted scRNA-seq on synoviocytes isolated from R805-treated and control animals. Unsupervised clustering identified several clusters expressing genes associated with macrophages (Fig. S26, **Table S2**); re-clustering of these cells defined 5 clusters (Fig. 6A). The largest cluster was biased toward control cells and was enriched for cells expressing anti-inflammatory macrophage markers including CD206/Mrc1 and CD163 (*55*) (Fig. 6A). Notably, one cluster populated entirely by cells derived from control animals expressed macrophage markers at lower levels but concurrently expressed high levels of the secreted collagenases MMP1 and MMP3 along with NOS2/Inos (Fig 6B). Moreover, this population also expressed COL1A1 and COL3A1, suggestive of a potential fibrocyte identity (*56*). Re-clustering of synovial fibroblasts defined a clear bias in clustering based on drug treatment (Fig. 6C**, ovals)**. Gene ontology analysis of genes significantly enriched in control vs. R805-treated fibroblasts (FDR <0.05, FC >1.5) revealed significant enrichment of categories related to inflammatory signaling and recruitment of inflammatory cells (Fig. 6C), anchored by enrichment of genes including *IL36B, PTGS2 (COX2), IL36G, SAA1, TLR4, CCL3* and *IL6* (**Table S3**). Together, these data suggest that R805 treatment biases the joint environment away from a chronic inflammatory identity following injury, therefore reducing the progression of cartilage degeneration and synovial fibrosis. Importantly, these findings were consistent with the data previously obtained from F814 mouse skin. Overall, the results from this study present a compelling case for a novel and potential therapeutic drug candidate for OA.

**Figure 6.**
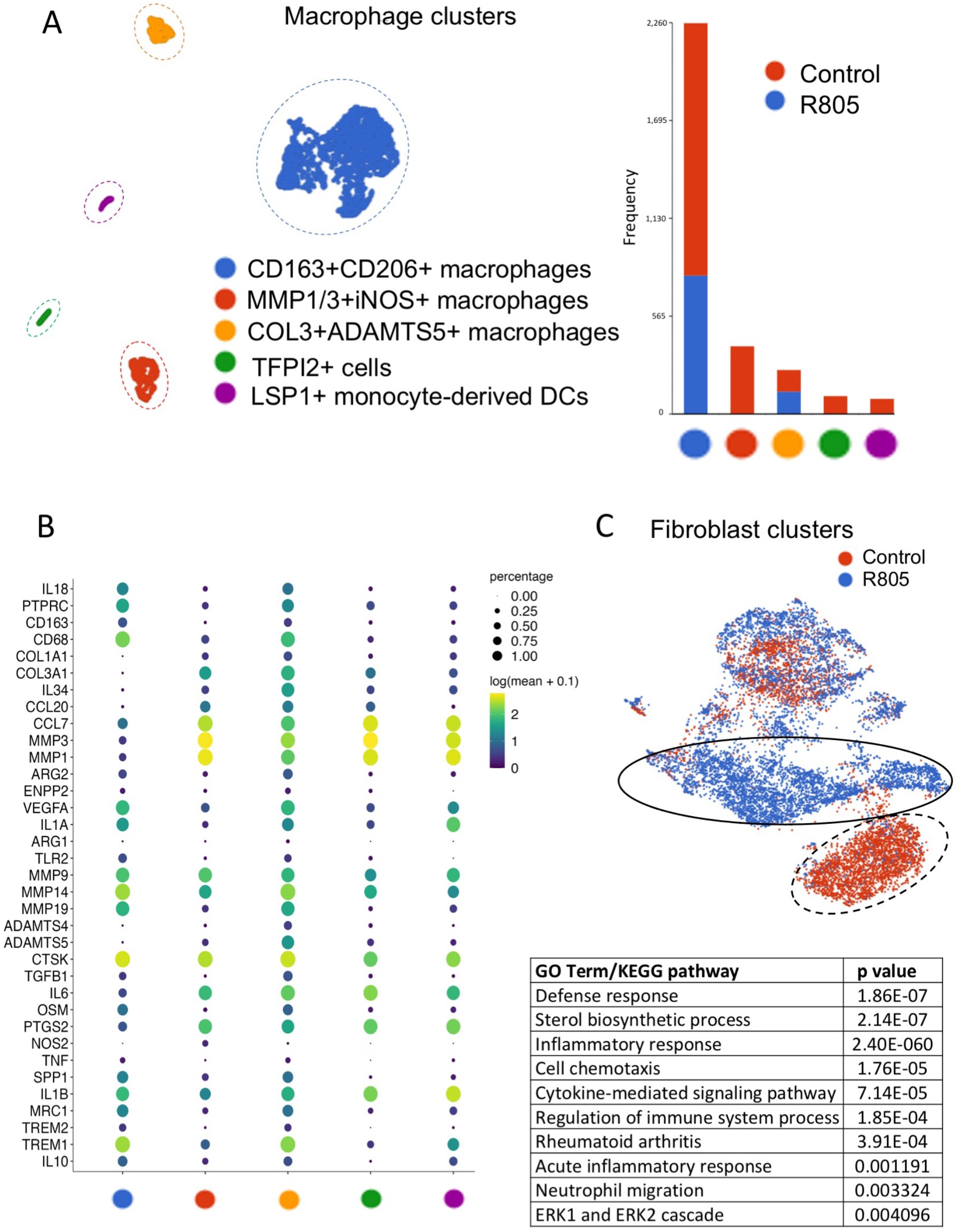
R805 promotes an anti-inflammatory, anti-fibrotic milieu in an injured joint. **A.** Re-clustering of clusters annotated as macrophages from synovial joint of R805-treated and vehicle-treated canine group. Clusters are named based on highly significant biomarkers and gene expression profiles. The contribution of each sample to each cluster is shown as a stacked bar graph. **B.** Dot plots depicting gene expression in each macrophage cluster from synovial joint of R805-treated and vehicle-treated canine group. Dot sizes are proportional to the percentage of cells in each cluster expressing the indicated gene. **C.** Re-clustering of clusters annotated as fibroblasts from synovial joint of R805-treated and vehicle-treated canine group. Cells are colored by their sample of origin. The dashed oval indicates clusters of fibroblasts dominated by R805-treated cells, while the solid oval denotes a cluster mainly derived from control cells. Select gene ontology terms over-represented in genes significantly enriched when comparing cells in the solid vs. dashed ovals.

## Discussion

The best characterized facet of gp130 signaling is its ability to promote or suppress inflammation resulting in tissue regeneration or pathology. This suggests that the divergence in outcomes downstream of gp130 are differentially regulated, which imparts the need for identification of a novel specific modality within this network that can be manipulated to initiate beneficial outcomes. Gp130 regulates signaling cascades via recruitment of proteins on its various residues; however, some of these signaling residues have not been well-characterized. We have demonstrated, for the first time, that a signaling Y814 residue within gp130 intracellular domain serves as a major cellular stress sensor that is responsible for inducing pro-inflammatory and pro-fibrotic outcomes along with SRC kinase recruitment. Instigation and maintenance of inflammation during disease pathogenesis is carried out by pro-inflammatory mediators that are tightly regulated. Our data presented here validates that Y814 is responsible for regulating most of the genes that are involved in inflammation and fibrosis including IL-6 cytokines, proteases, cellular adhesion molecules that are critical in leukocyte recirculation, COX2 and others. Since constitutive inflammation is the foremost culprit in the dysregulation of normal wound healing, targeting Y814 may potentially orchestrate efficacious tissue regeneration processes.

Our previous studies have demonstrated that gp130-STAT3 signaling is highly upregulated in proliferative, anabolic fetal chondrocytes (*32*), and that LIF is highly expressed in developing human joints (*17*). Our most recent study demonstrated that Lifr-gp130-Stat3 signaling is required for homeostatic maintenance of chondroprogenitors in mice, and that genetic postnatal ablation of any members of this triad results in premature growth plate fusion and progressive changes in articular cartilage (*57*). Strikingly, genetic overexpression of STAT3 in postnatal chondrocytes did not induce an OA phenotype; moreover, chondrocyte hyperproliferation was observed in both growth plate and articular cartilage (*57*). Together, these data challenged previous oversimplified views on gp130-STAT3 signaling in cartilage tissue suggesting that activation of this pathway in OA may initially represent a regenerative attempt, but prolonged and excessive activation of this mechanism is likely to be detrimental. Current study demonstrates that genetic or pharmacological inhibition of gp130-Y814 module designed to minimally interfere with the endogenous STAT3 signaling significantly improves regenerative outcomes in multiple tissues.

Even though we have shown that gp130 Y814 serves as a recruitment site for SRC kinase, it is still unknown whether Y814 is an exclusive recruitment site for solely SRC as SFK family consists of 9 family members that are neither universally expressed nor conserved (*58*). It is also difficult to distinguish among SFKs as current available reagents lack complete discrimination to discern the kinases completely. Mutation in Y814 also downregulates MAPK (ERK 1/2) signaling, which is a central pathway controlling cellular processes associated with fibrogenesis, including growth, proliferation, and survival (*59, 60*). It is unclear whether SRC and MAPK/ERK crosstalk on Y814, but is conceivable as SRC is known to activate ERK signaling in multiple pathologies (*61, 62*). Conversely, other pathways downstream of gp130 such as STAT3, YAP1 and AKT remained minimally affected.

Since our RNA seq analysis demonstrated that ablation of Y814 *in vivo* can dramatically reduce expression of multiple pro-inflammatory and pro-fibrotic genes in response to OSM (e.g. *Cox2*, *Il-11*, *Nf-kb*, etc.), it was not unanticipated that in the skin wound healing and joint destabilization injury models, Y814 transgenic mice (F814) demonstrated a regenerative, anti-fibrotic healing potential while averting degenerative processes, respectively. It is well known that a considerable number of factors contribute to decline in regenerative capability and susceptibility to degeneration (*1*). Based on our data, activation of this pro-inflammatory and pro-fibrotic stress sensor may be one of the factors limiting regenerative capacity. Genetic knockout of Y814 resulted in a decrease of inflammatory gene signature as visualized by the anti-inflammatory macrophage and non-pathological fibroblast subpopulations in the damaged skin tissue, and this improvement in the microenvironment may contribute to the observed enhanced regeneration.

In addition, our fundamental studies in the areas of chronic inflammation have identified a novel class of small molecules capable of selective manipulation of gp130 receptor (*32*). Here we show that an optimized RCGD 423 analog, R805, prevents Y814 activation in response to IL-6 cytokines conveying specific mode of regulation without hindering other pro-regenerative pathways and dramatically reduces the appearance of OA in rat and canine models. Similar to the ablation of the Y814 in mouse, R805 decreases the pathological milieu in mouse skin wound and diseased canine joint. In combination with the lizard tail regeneration, all these novel findings suggest that both Y814 and, consequently, R805 may modulate beneficial outcomes via macrophage polarization towards an anti-inflammatory phenotype; however, further elucidation is necessary to confirm which cell populations are involved, and this is beyond the scope of the current study.

Thus, in this study, we present a specific, biologically essential Y814 cellular stress sensor within gp130 receptor, which serves as a novel therapeutic target for drug development and holds considerable promise from a translational standpoint. Pharmacological manipulation of specific gp130 modules that enhance intrinsic regenerative capacity of tissues and resistance to degenerative changes can offer safe symptomatic treatments and alternatives to the conventional treatment options that could revolutionize the contemporary approach for a broad range of conditions. Therapeutic approaches designed towards enhancement of regenerative abilities could also result in substantial lifespan improvements.

## Acknowledgments

**General:** De-identified human cartilage samples were collected under UCLA IRB# 10-001857. We would like to thank Ms. Jade Tassey (USC) for preparing single cell suspension of canine synovial samples for single cell analysis.

## Author contributions

Conceptualization: DE, RS, CMC, BVH, MK

Methodology: RS, CF, EL, AS, JL, SL, JL, HICH, KV, PB

Investigation: SBB, DLA, MPW, WCB, KV, PB

Visualization: HICH, NQL, MH

Funding acquisition: DE

Project administration: DE, MH, CMC

Supervision: DE, NQL, MH, CMC

Writing – original draft: RS, DE, MH, CMC

Writing – review & editing: RS, DE, MH, BVH, CMC, AS

## Data and materials availability

All RNA sequencing data for this manuscript will be available via a publicly open database GEO (See Materials and Methods). Genetic F814 mouse and polyclonal antibodies for Y814 residue will be available to academic researchers via materials transfer agreements (MTAs).

---

**Fig. S1.**
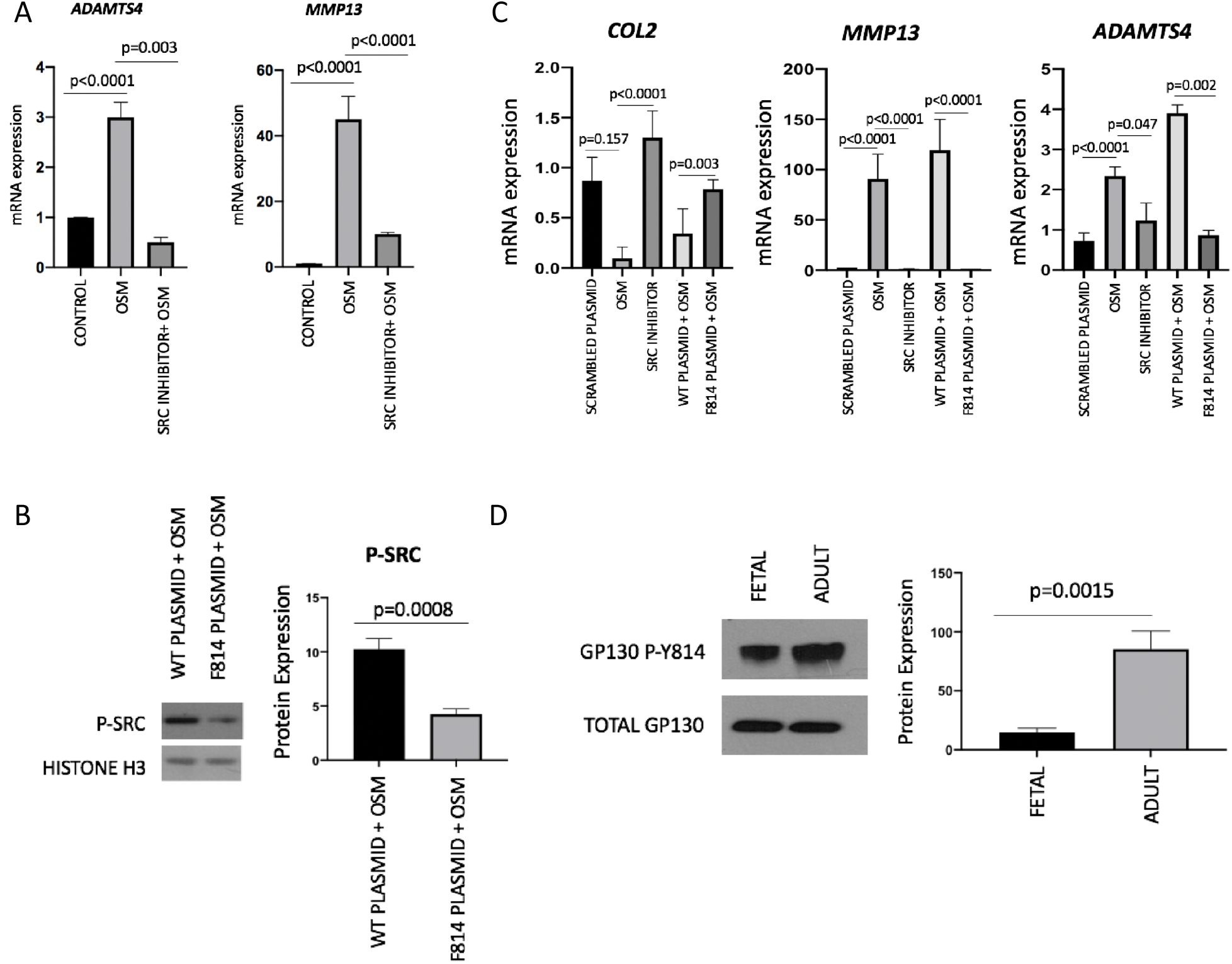
Y814 residue regulates SRC signaling downstream of gp130. **A.** qPCR results from pig articular chondrocytes stimulated with OSM and treated with or without SRC inhibitor (SU6656) for 48 hours. Horizontal lines with bars show the mean ± SD. n=3. **B.** Protein expression of pSRC in ATDC5 cells after transfection (72 hours) with the modified and WT gp130 plasmids stimulated with OSM for 24 hours. Horizontal lines with bars show the mean ± SD. n=3. **C.** qPCR results from ATDC5 cells treated with SRC inhibitor (SU6656) alone, or transfected with indicated variants of plasmids and stimulated with or without OSM for 48 hours. Horizontal lines with bars show the mean ± SD. n=3. **D.** Protein expression of gp130 pY814 in untreated fetal (17 weeks) and adult articular chondrocytes. Horizontal lines with bars show the mean ± SD. n=3.

**Fig. S2.**
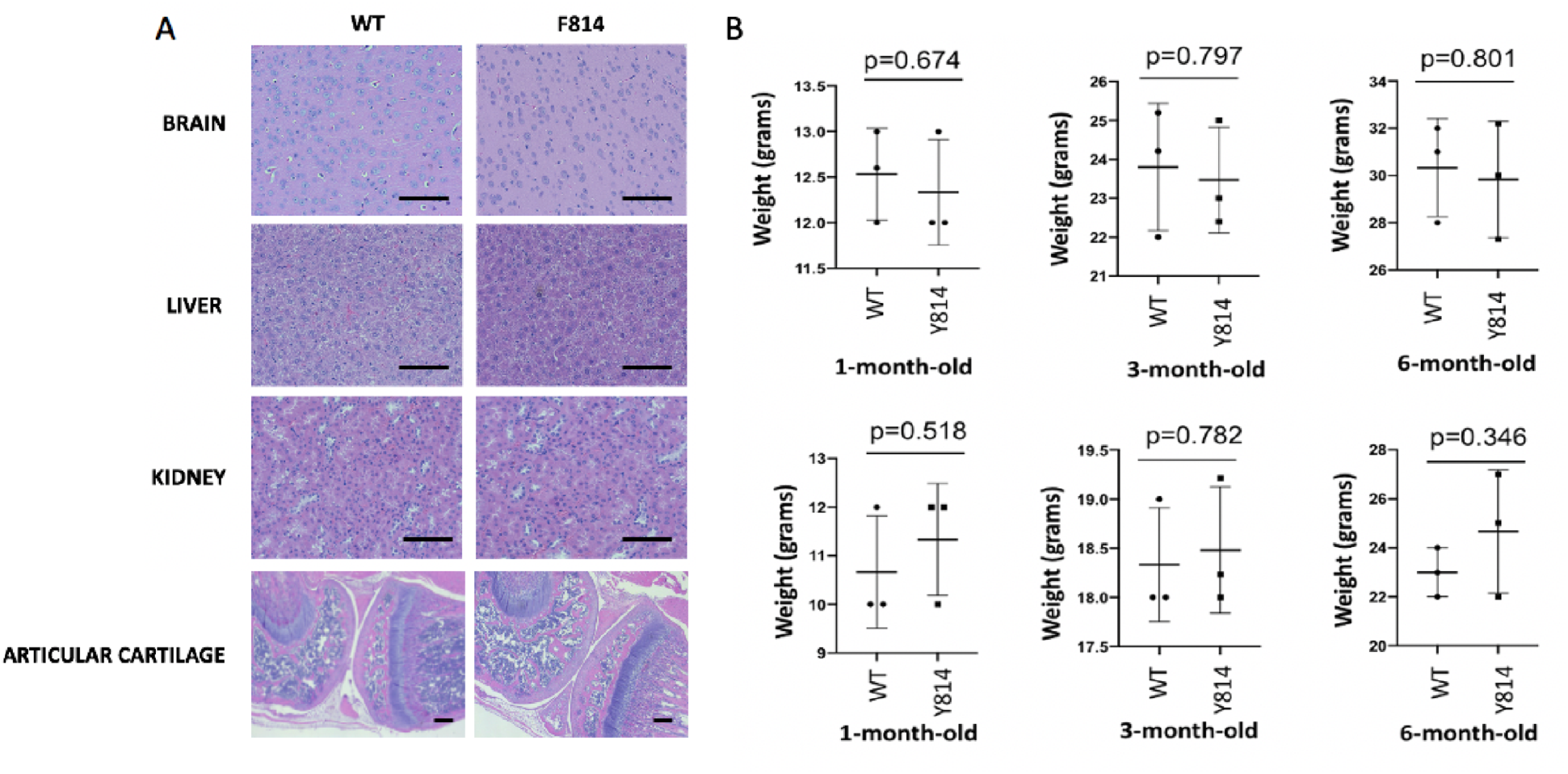
F814 mouse exhibits no noticeable morphological differences in major organs. **A.** H&E staining for indicated wild type (WT) and F814 mouse organs. Magnification=20X. n=3. **B.** Graphs indicate the average body weight (grams) of 4-month-old males. Representative images are shown. Horizontal lines with bars show the mean ± SD. n=3.

**Fig. S3.**
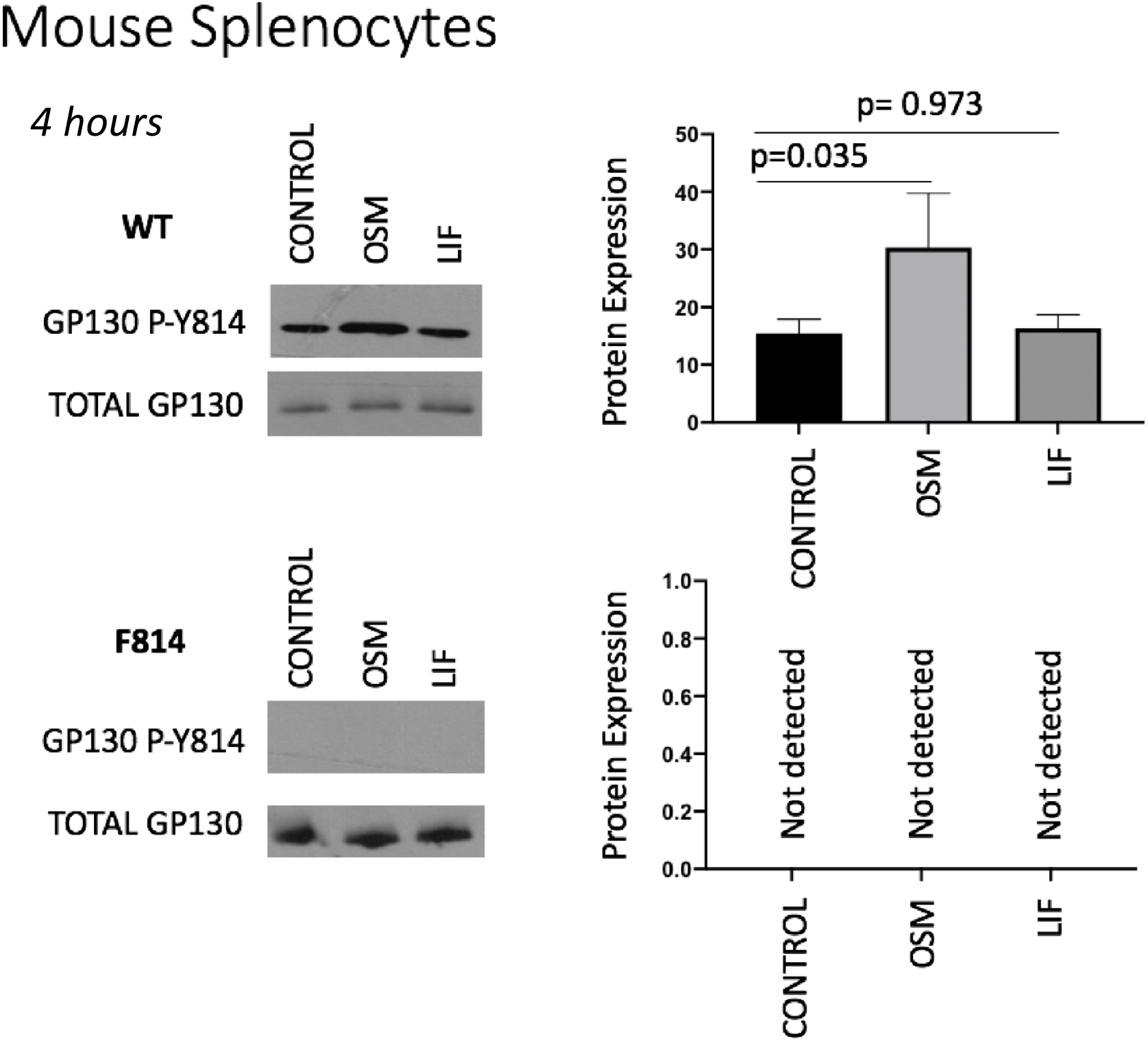
F814 mouse does not express gp130 Y814. Protein expression of gp130 pY814 in wild type (WT) and F814 mouse splenocytes treated with and without OSM and LIF for 4 hours. Horizontal lines with bars show the mean ± SD. n=3.

**Fig. S4.**
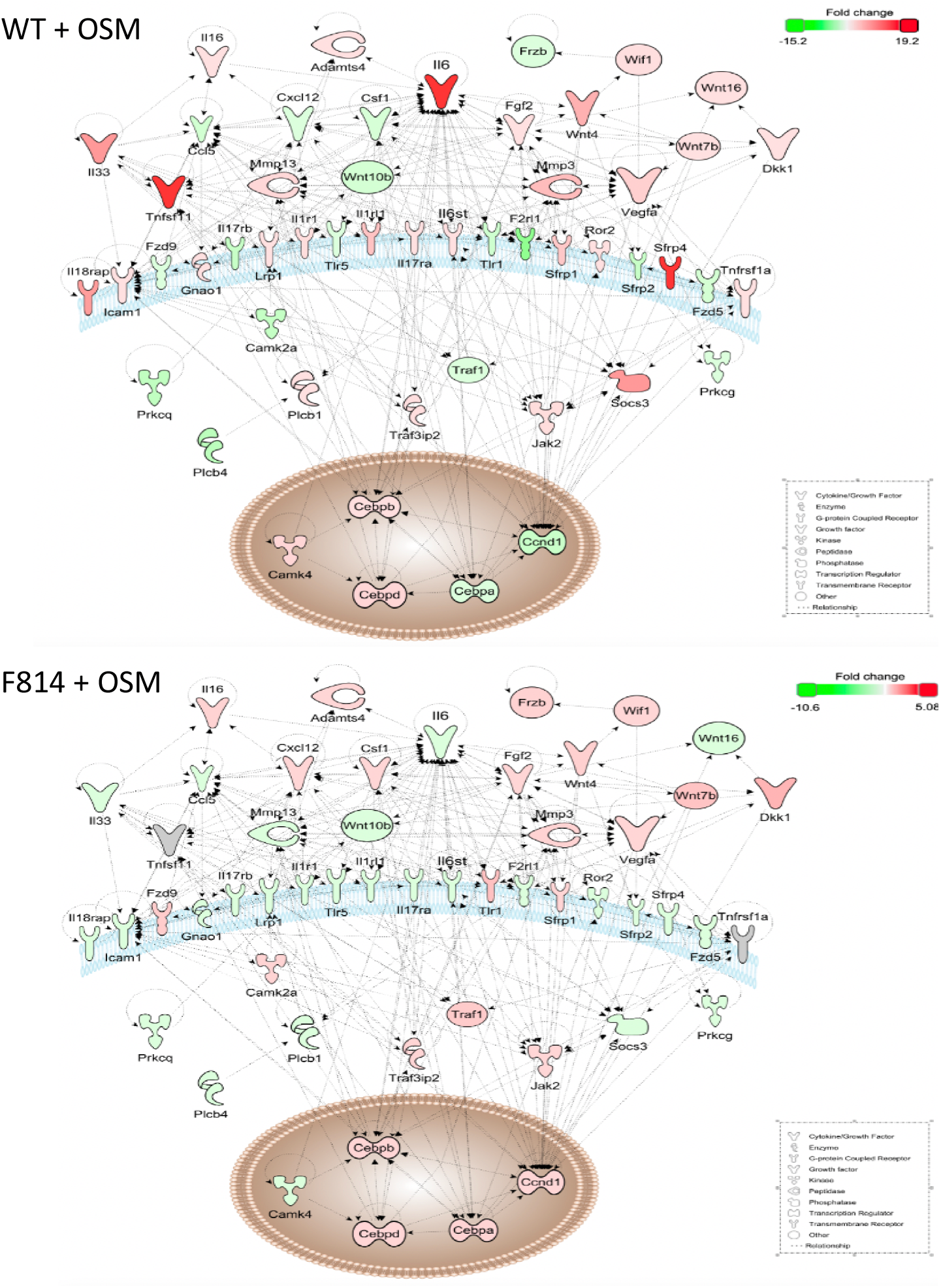
OA Ingenuity pathway analysis. RNA-seq results demonstrated that deletion of Y814 can dramatically reduce expression of pro-inflammatory genes in response to OSM with Ingenuity Macrophage RA pathway being one of the most affected by this mutation. Murine primary fibroblasts were stimulated with OSM for 24 hours.

**Fig. S5.**
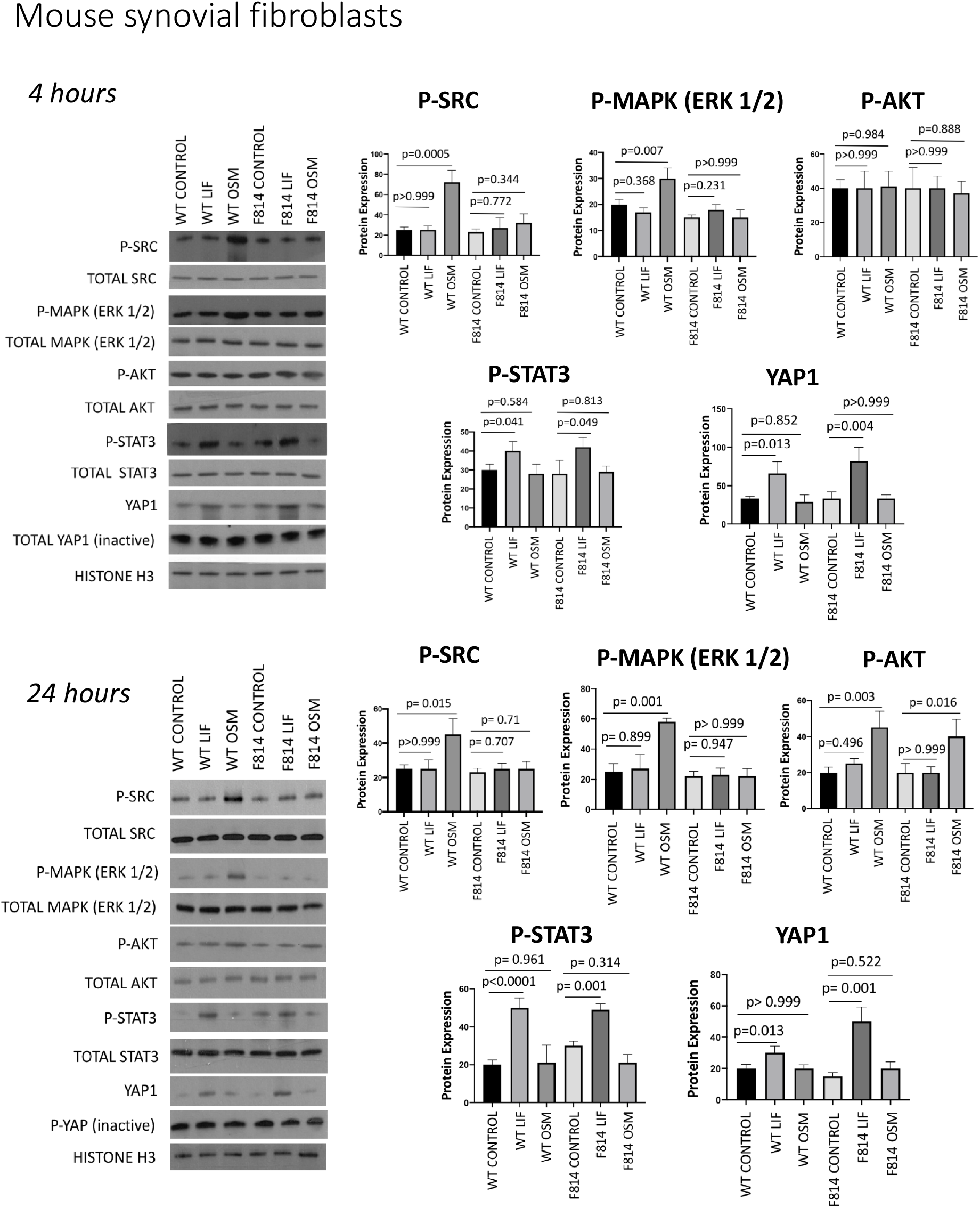
Wild type and F814 mice have different modes of regulation downstream of gp130. Protein expression of indicated proteins in wild type (WT) and F814 mouse primary periarticular fibroblasts stimulated with or without OSM or LIF for 4 and 24 hours. Horizontal lines with bars show the mean ± SD. n=3.

**Fig. S6.**
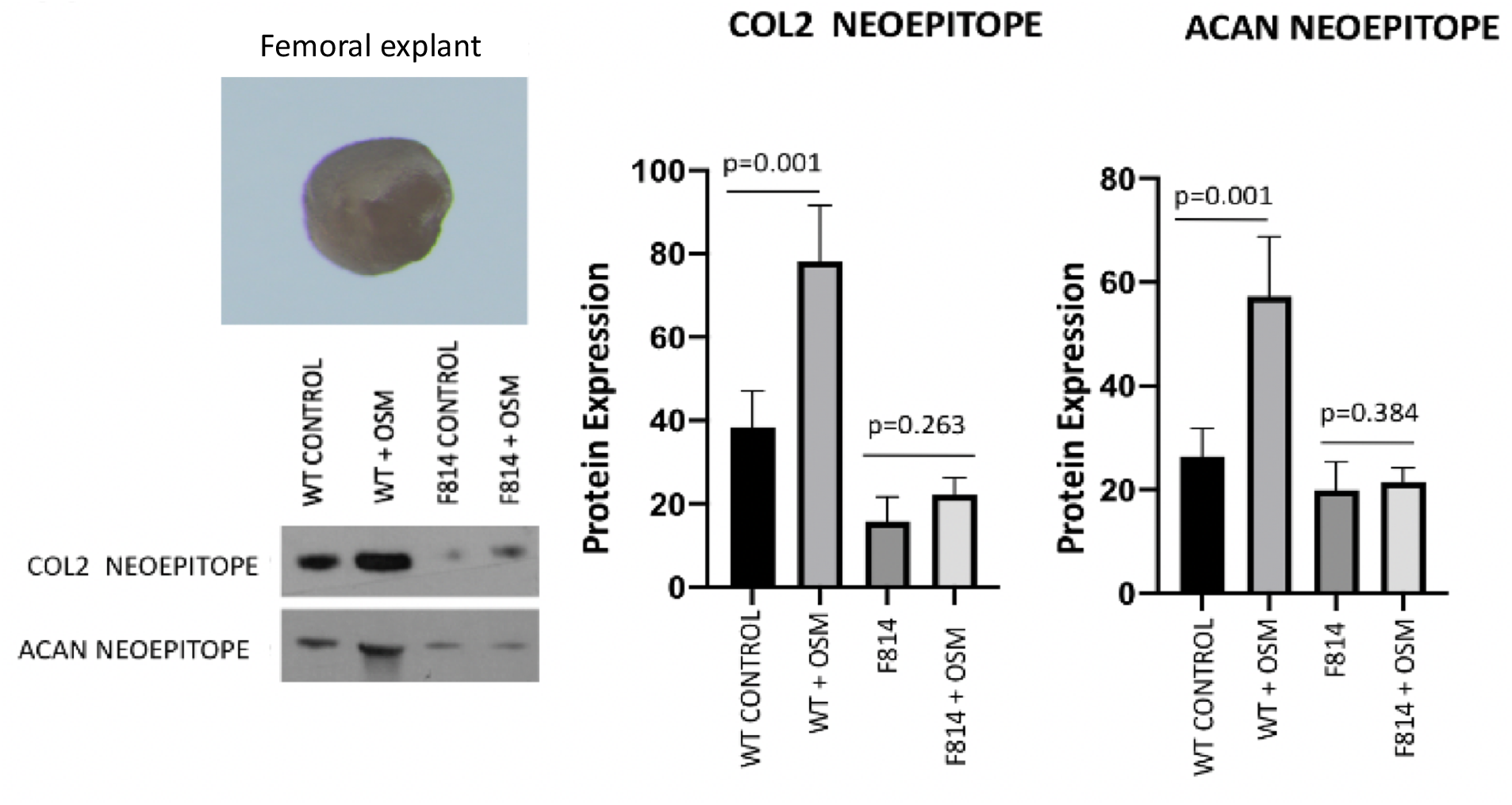
Ablation of Y814 inhibits OSM-induced collagenase and aggrecanase activity. Wild type and F814 mouse knee articular cartilage explants were incubated with or without OSM for 72 hours followed by a neoepitope assay. Horizontal lines with bars show the mean ± SD. n=3.

**Fig. S7.**
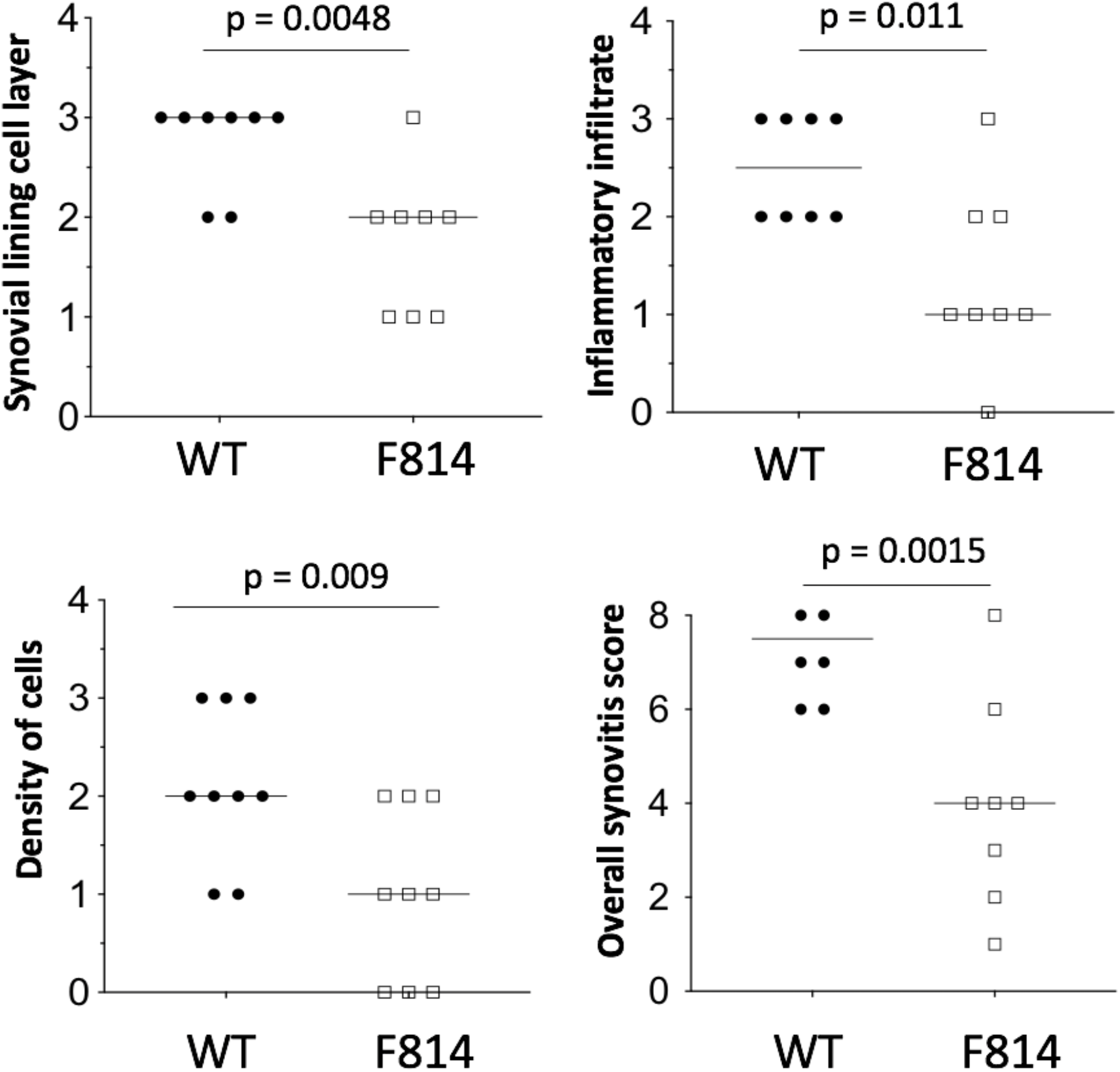
F814 mouse exhibits reduced synovitis after surgically induced osteoarthritis. Quantitative assessment of synovial lining cell layer, inflammatory infiltrate and density of cells in wild type (WT) and F814 in mouse knee joints 6 weeks after destabilization of the medial meniscus surgery. n=9.

**Fig. S8.**
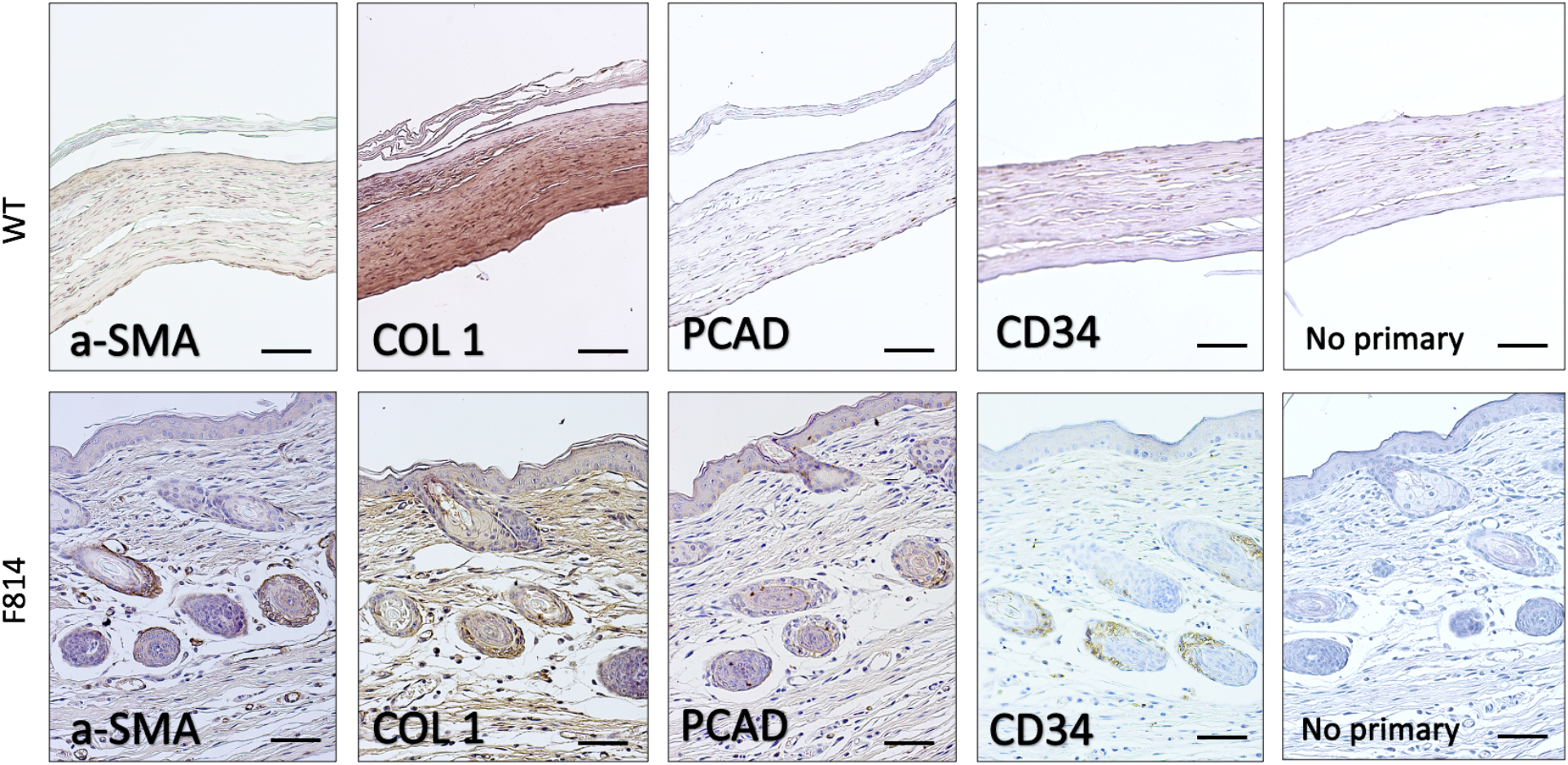
F814 mouse exhibits enhanced wound-induced hair neogenesis. Wild type (WT) and F814 mouse post-wound day (PWD) 21 wound sections after wound excision. Representative images are shown. n=6.

**Fig. S9.**
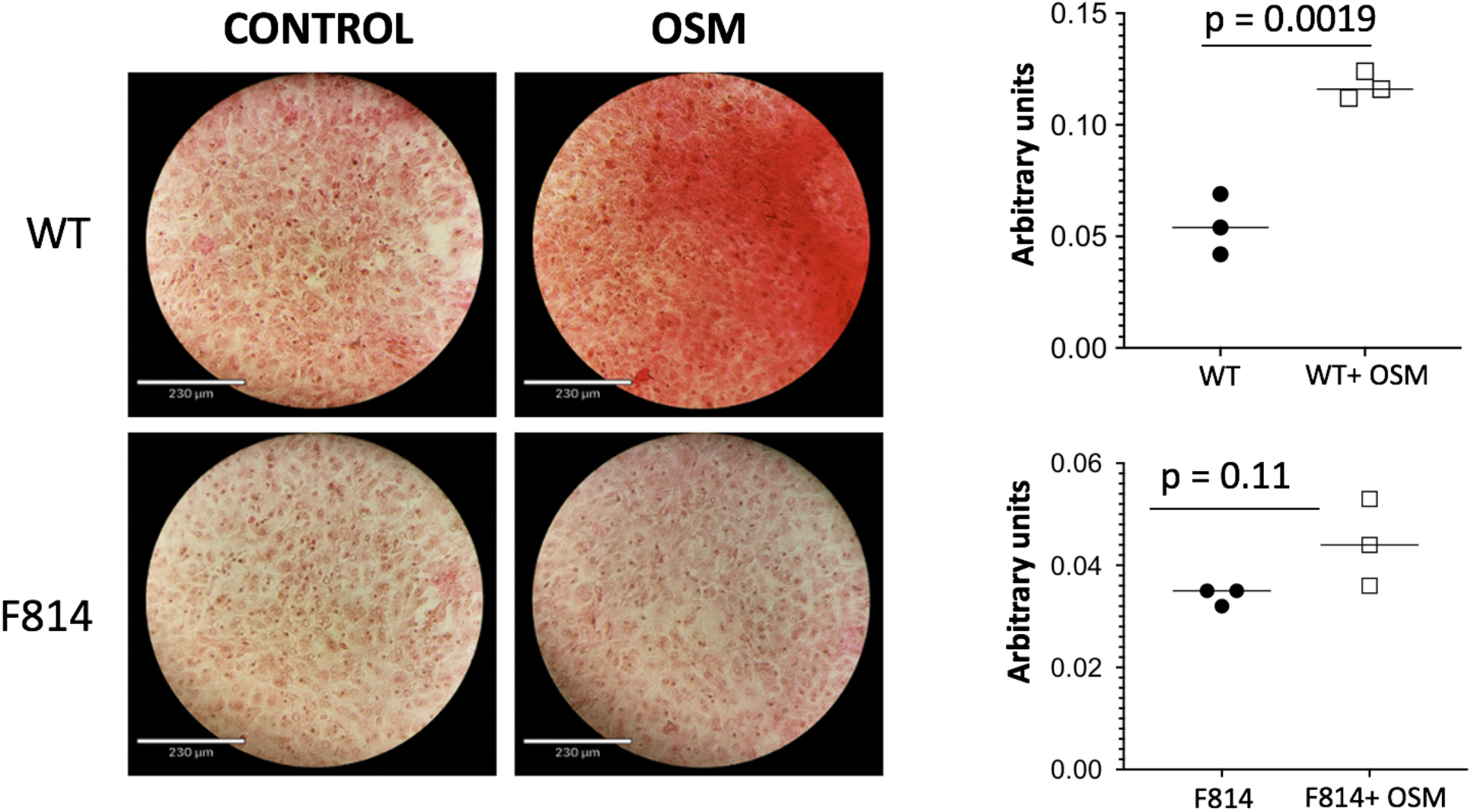
F814 mouse cells are resistant to OSM-induced collagen deposition. Picrosirius Red staining of periarticular subcutaneous adipose fibroblasts in wild type (WT) and F814 mouse. Representative images are shown. n=3.

**Fig. S10.**
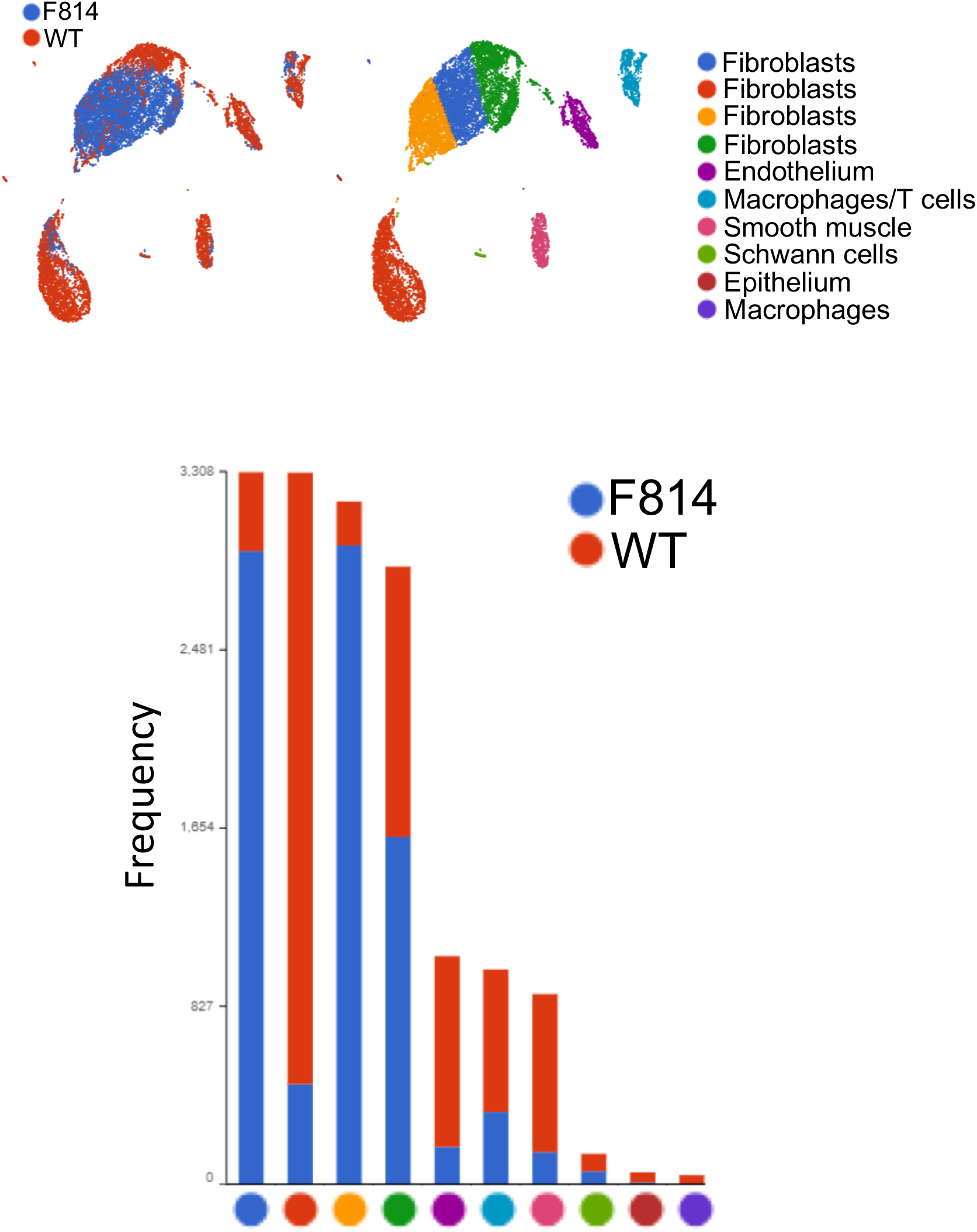
scRNA-sequencing of large wounds from F814 and wild type (WT) mice. UMAP and k-means clustering of equal numbers of cells isolated at post-wound day (PWD) 14 from the indicated genotype; the number of cells in each cluster from each genotype is displayed in the stacked bar graph. Presumptive identities of each cluster are displayed based on biomarker genes significantly enriched; see also Supp. Table X. Cells were harvested from wounds pooled from 2-3 animals of each genotype and sorted as DAPI and Ter119 negative then subjected to the 10X Genomics workflow.

**Fig. S11.**
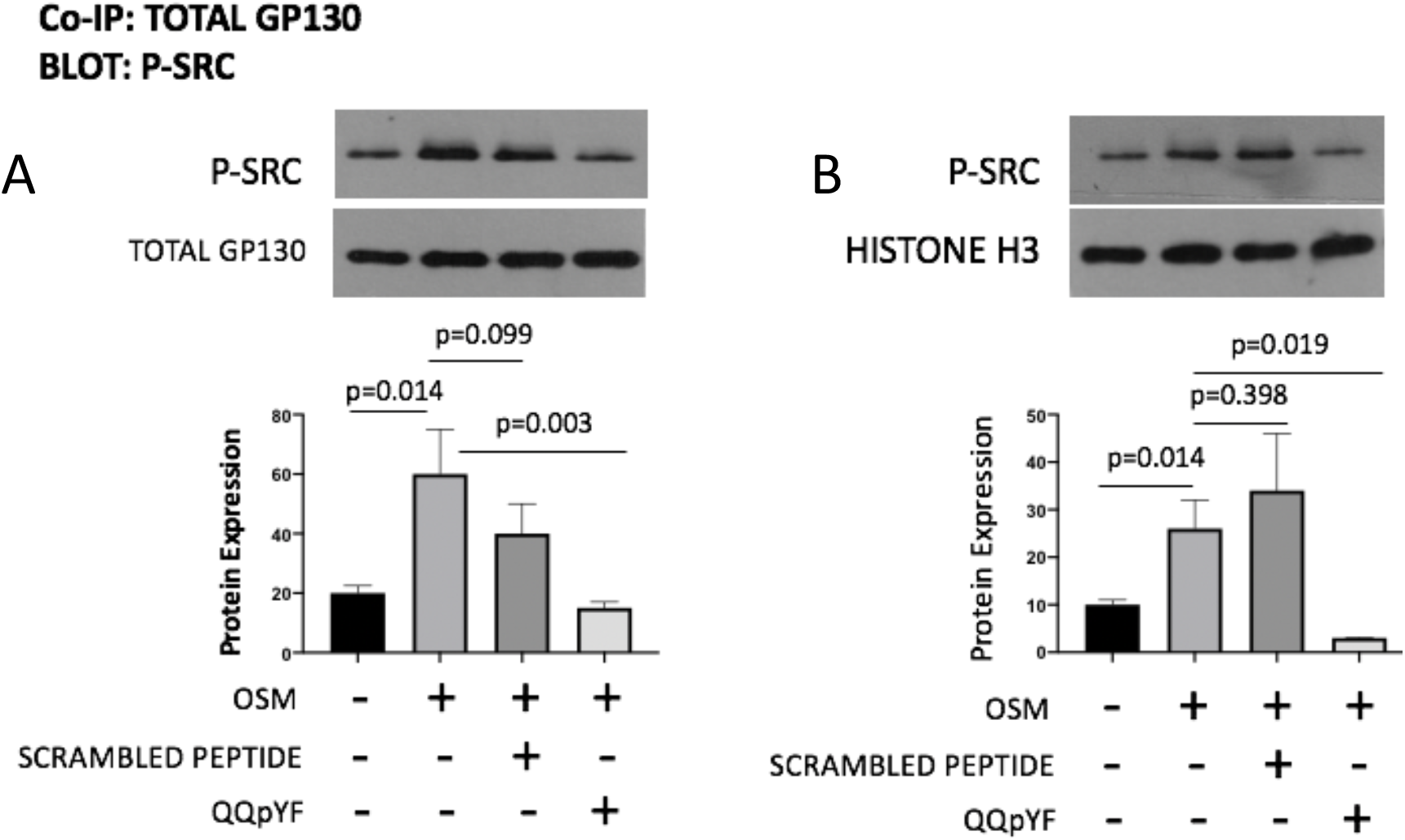
Peptide QQpYF inhibits activation of SRC downstream of gp130. **A.** Levels of complex formation between gp130 and pSRC in pig articular chondrocytes stimulated with or without OSM in presence or absence of control scrambled peptide or peptide QQpYF for 4 hours. Horizontal lines with bars show the mean ± SD. n=3. **B.** Protein levels of pSRC in pig articular chondrocytes stimulated with or without OSM in presence or absence of control scrambled peptide or peptide QQpYF for 4 hours. Horizontal lines with bars show the mean ± SD. n=3.

**Fig. S12.**
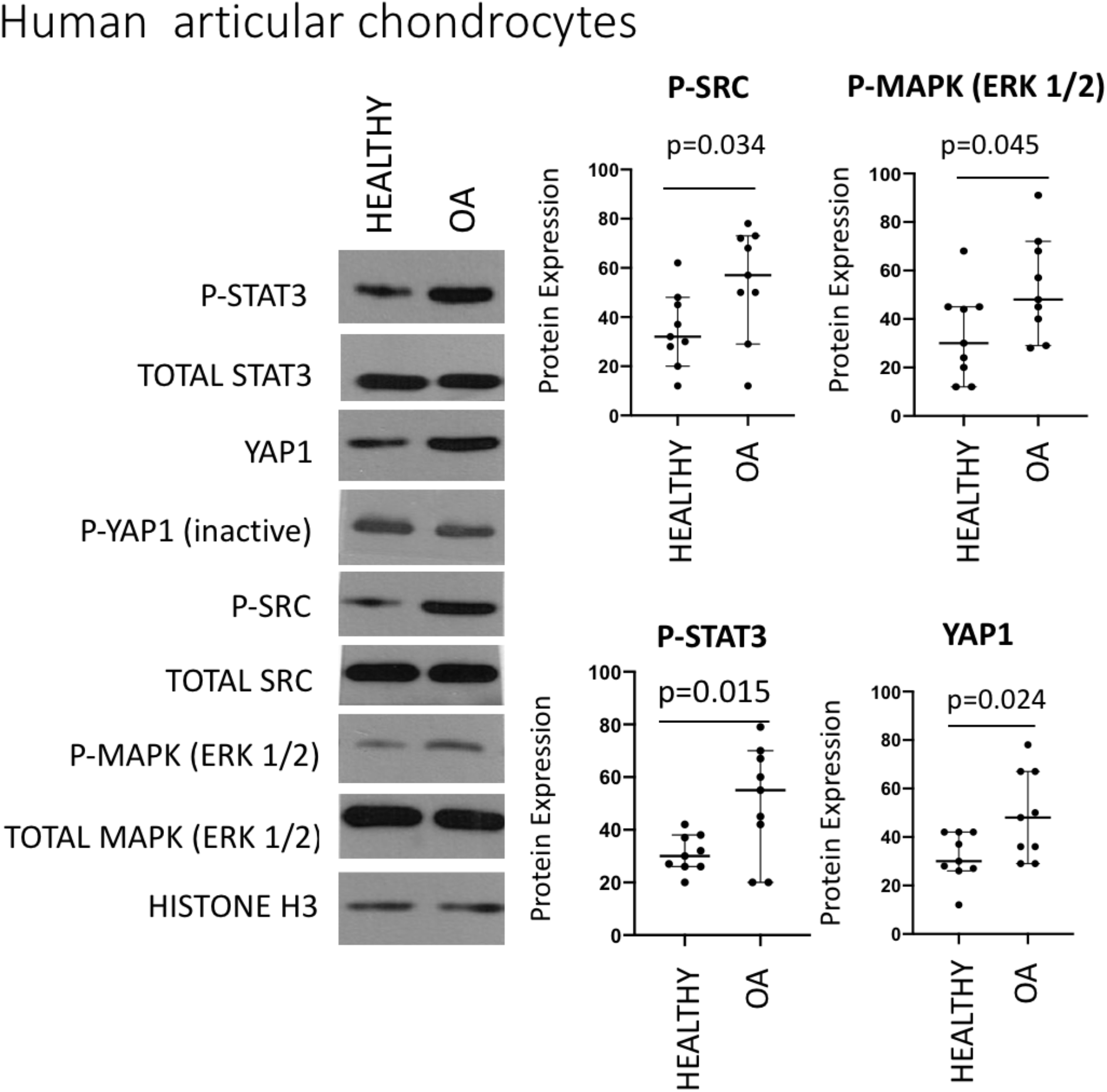
STAT3 upregulation is high in diseased tissue. Endogenous protein expression of indicated proteins in healthy and osteoarthritic chondrocytes. Horizontal lines with bars show the mean ± SD. n=9.

**Fig. S13.**
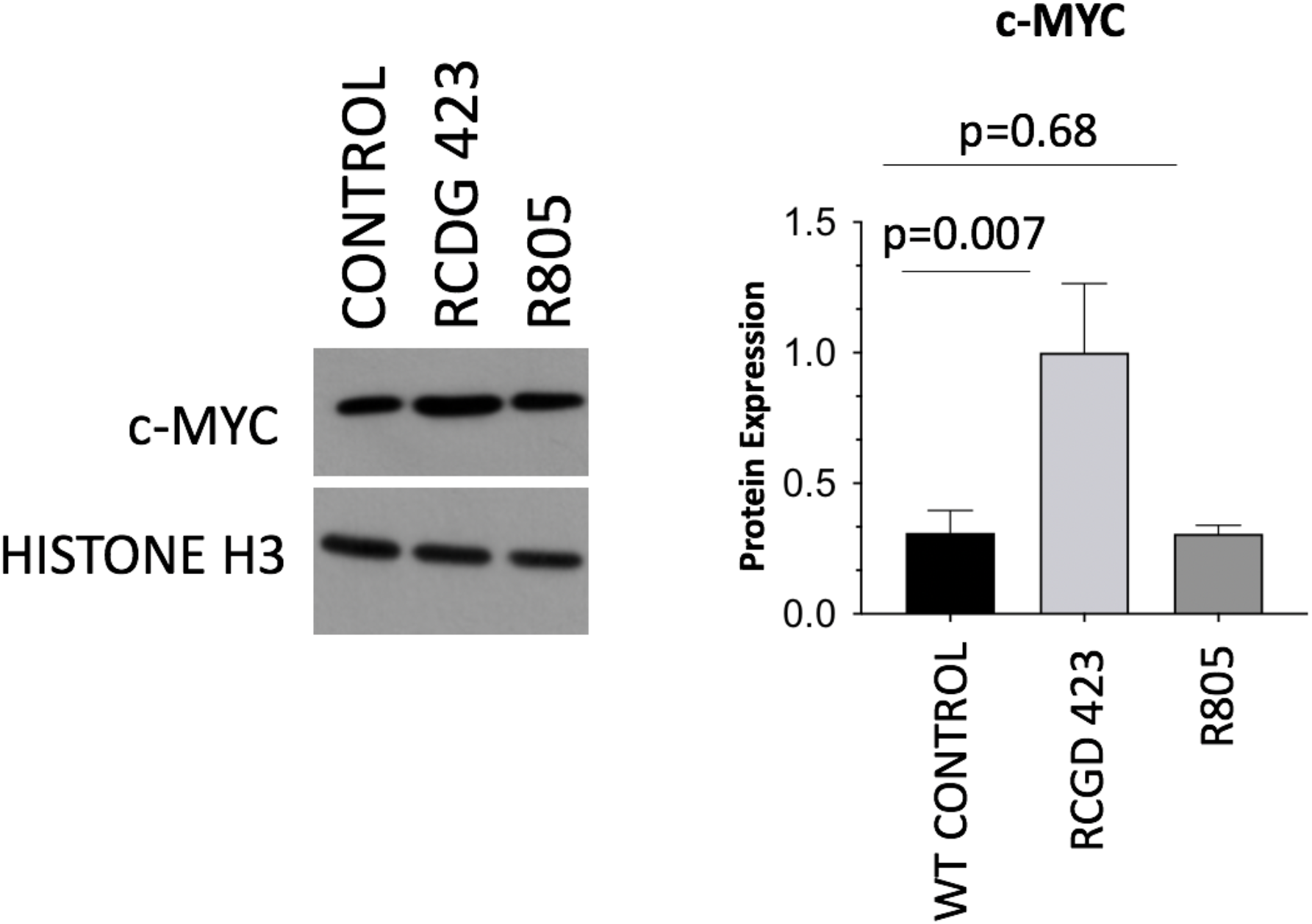
R805 does not activate MYC signaling. Protein levels of c-MYC in pig articular chondrocytes stimulated with or without RCGD423 or R805. Horizontal lines with bars show the mean ± SD. n=3.

**Fig. S14.**
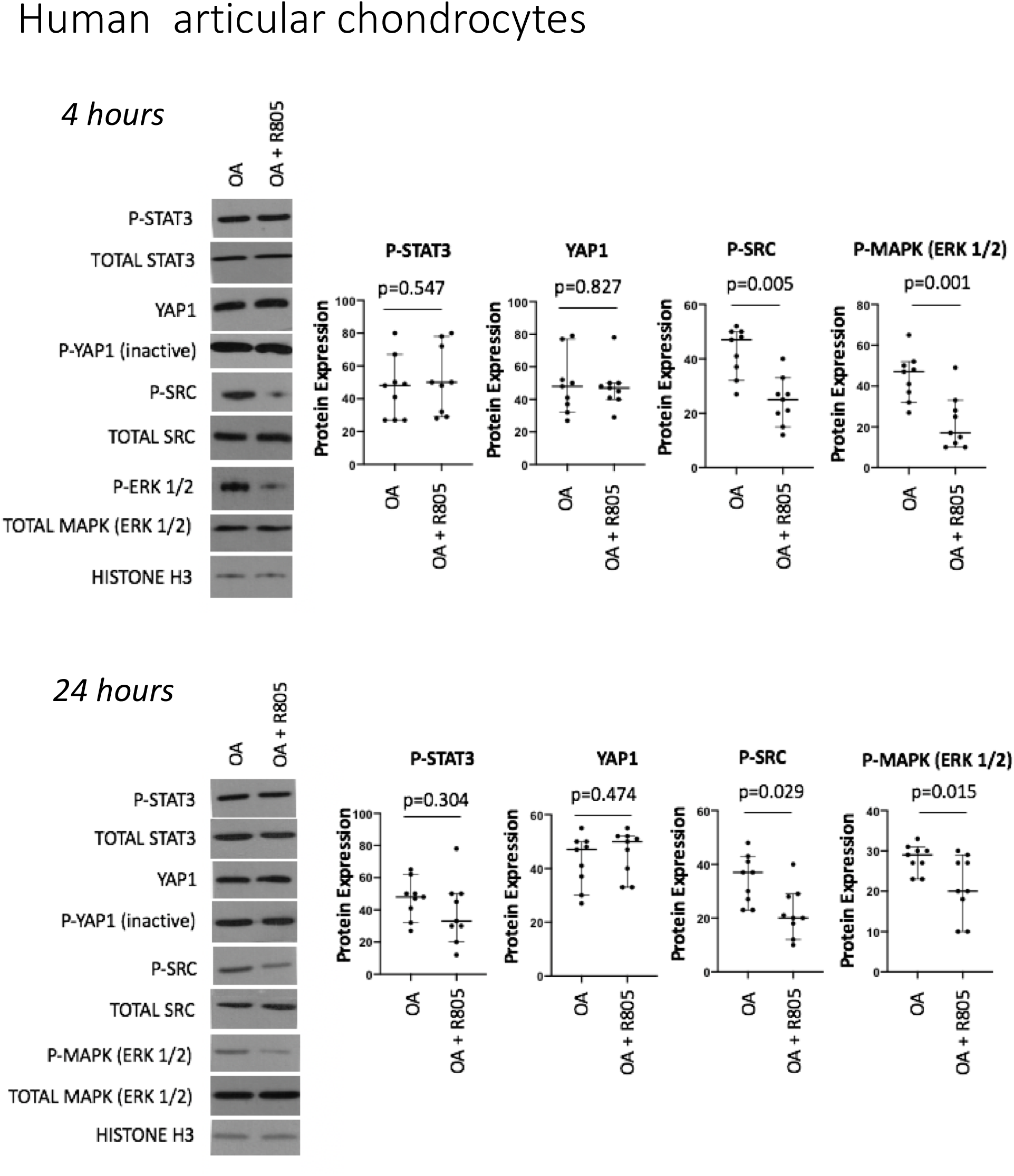
R805 prevents pro-inflammatory signaling in diseased tissue. Protein expression of indicated proteins in osteoarthritic chondrocytes treated with or without R805 for 4 and 24 hours. Protein levels were quantified with respect to Histone H3. Horizontal lines with bars show the mean ± SD. n=9.

**Fig. S15.**
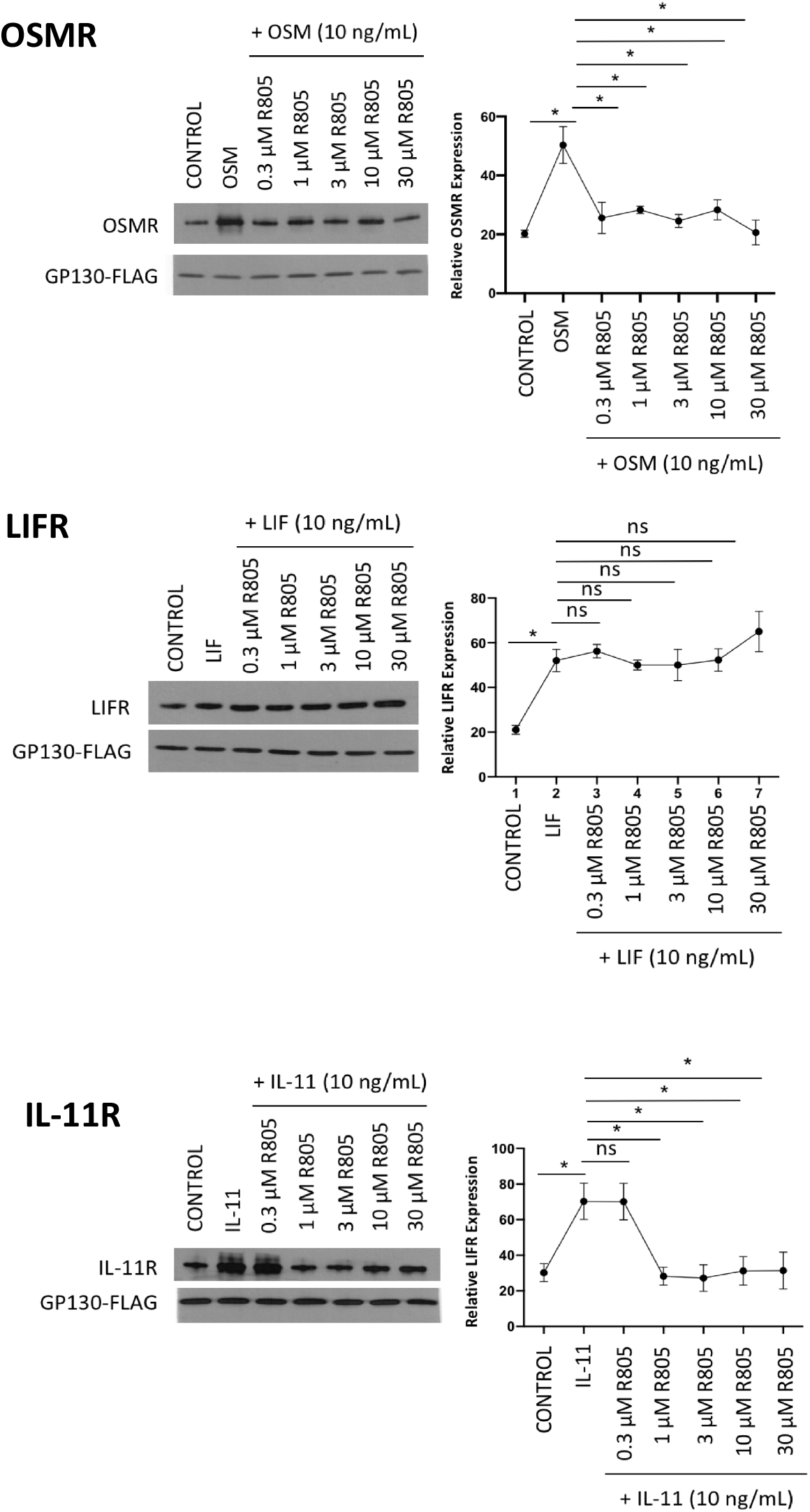
R805 inhibits heterodimerization of gp130 with IL-6 family of cytokines’ alpha receptors. Receptor-competition assay. Adult human articular chondrocytes were treated with or without indicated IL-6 family of cytokines and different doses of R805 for 4 hours. Co-IP measuring protein expression was performed after transfection with gp130 FLAG plasmid (72 hours) followed by western blot with respective receptor antibodies. Horizontal lines with bars show the mean ± SD. n=3.

**Fig. S16.**
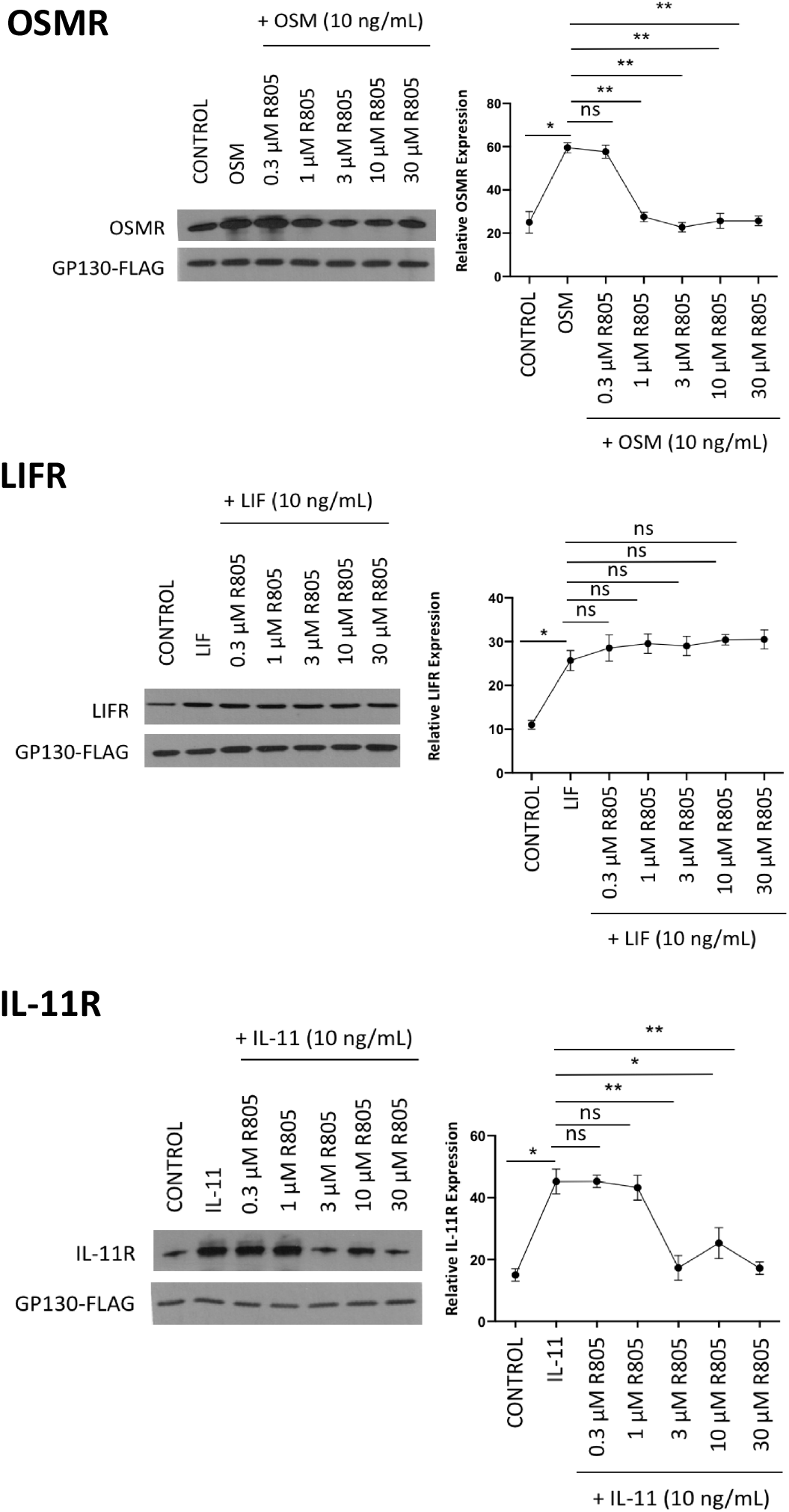
R805 inhibits heterodimerization of gp130 with IL-6 family of cytokines’ alpha receptors. Receptor-competition assay. Adult human articular chondrocytes were treated with or without indicated IL-6 family of cytokines and different doses of R805 for 24 hours. Co-IP measuring protein expression was performed after transfection with gp130 FLAG plasmid (72 hours) followed by western blot with respective receptor antibodies. Horizontal lines with bars show the mean ± SD. n=3.

**Fig. S17.**
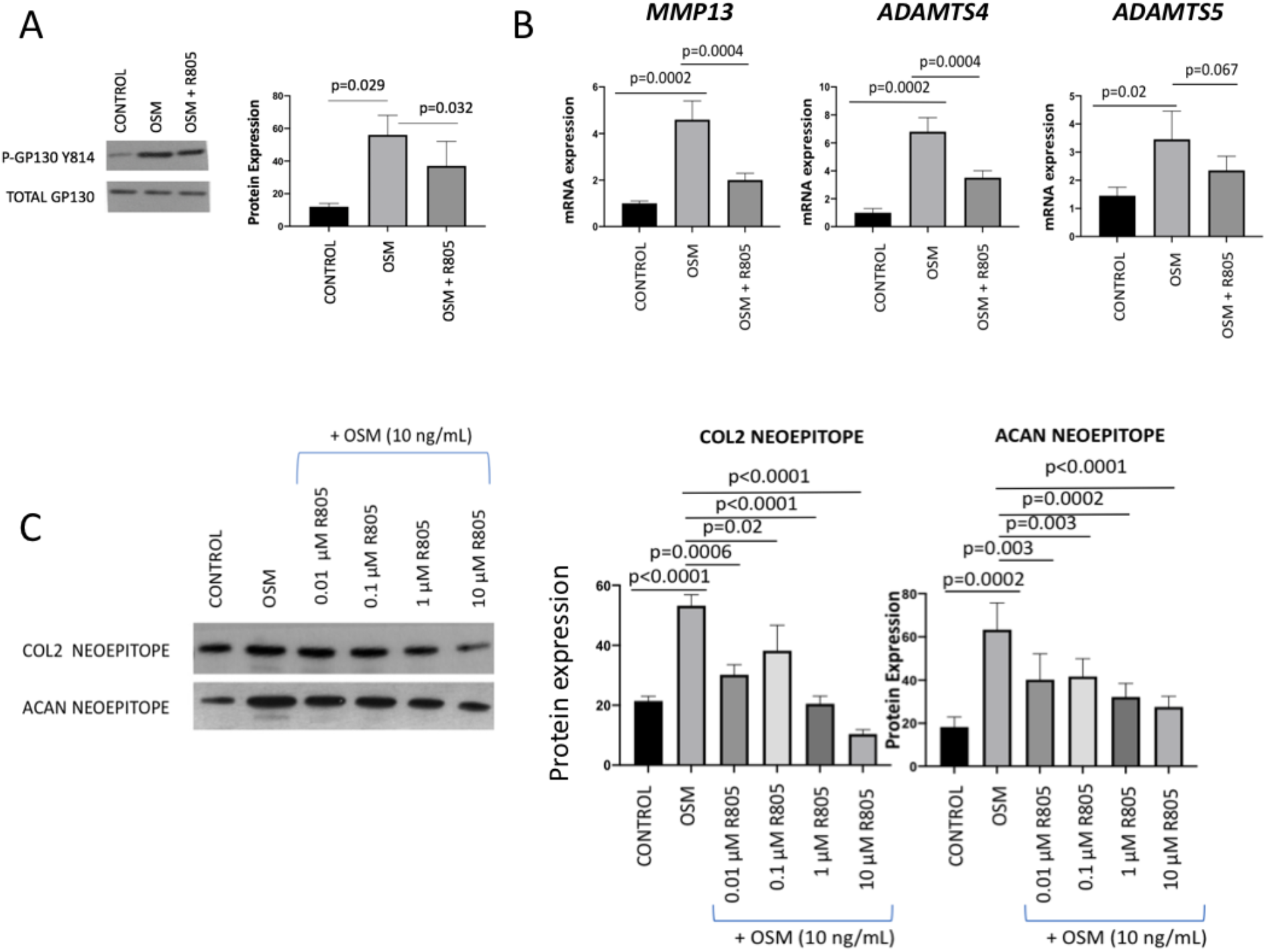
R805 exhibits a protective effect on OSM-induced cartilage degeneration *in vitro*. **A.** Protein expression of gp130 pY814 in pig articular chondrocytes treated with or without OSM and R805 for 4 hours. Horizontal lines with bars show the mean ± SD. n=3. **B.** qPCR results from pig articular chondrocytes cells treated with OSM with or without R805 for 48 hours. Horizontal lines with bars show the mean ± SD. n=3. **C.** Pig cartilage knee explants were incubated with or without OSM for 72 hours with or without R805 at indicated concentrations followed by a neoepitope assay. Horizontal lines with bars show the mean ± SD. n=3.

**Fig. S18.**
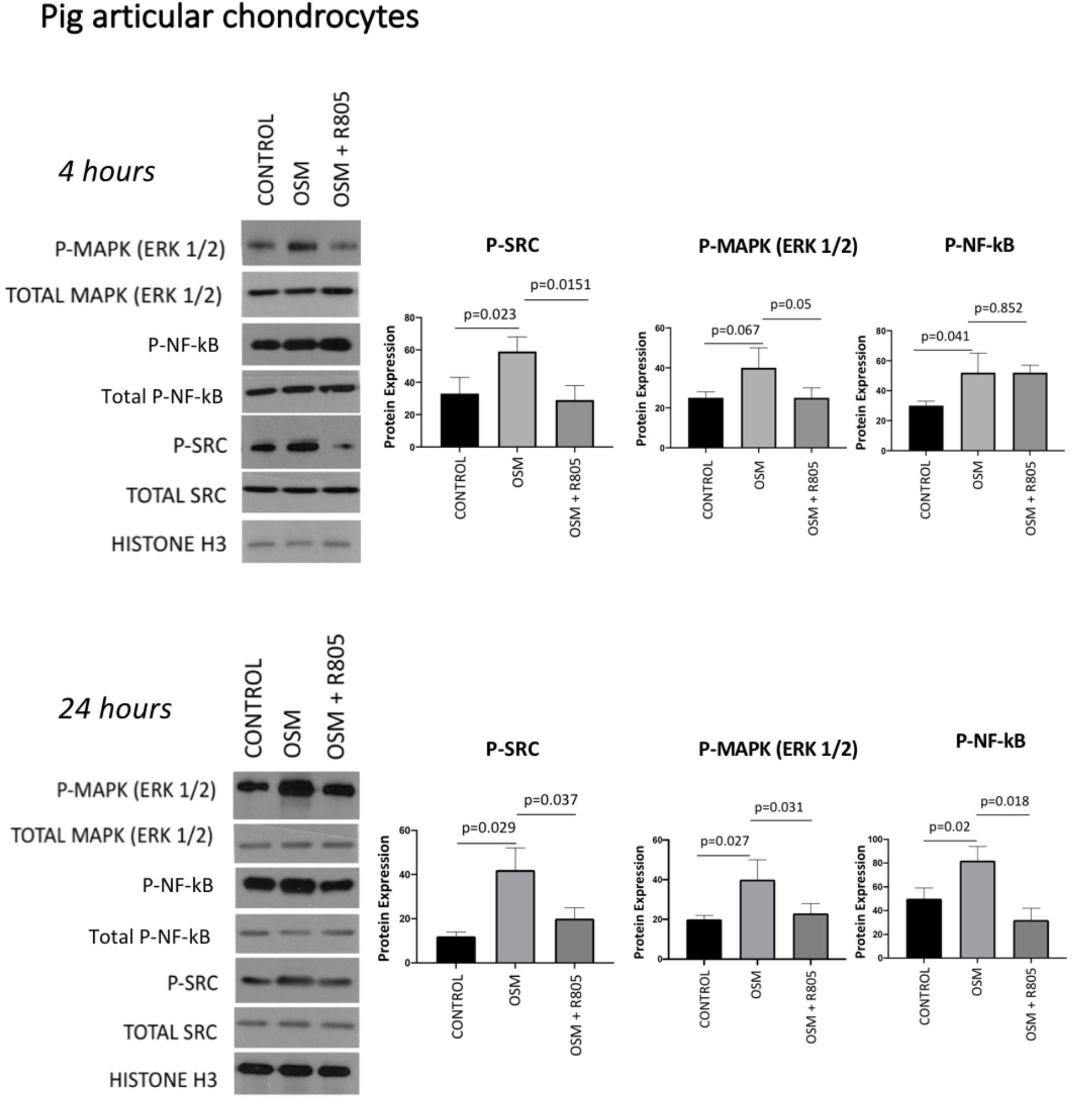
R805 prevents pro-inflammatory signaling in OSM-stimulated pig articular chondrocytes. Protein expression of pSRC in pig articular chondrocytes treated with or without OSM and R805 for 4 and 24 hours. Horizontal lines with bars show the mean ± SD. n=3.

**Fig. S19.**
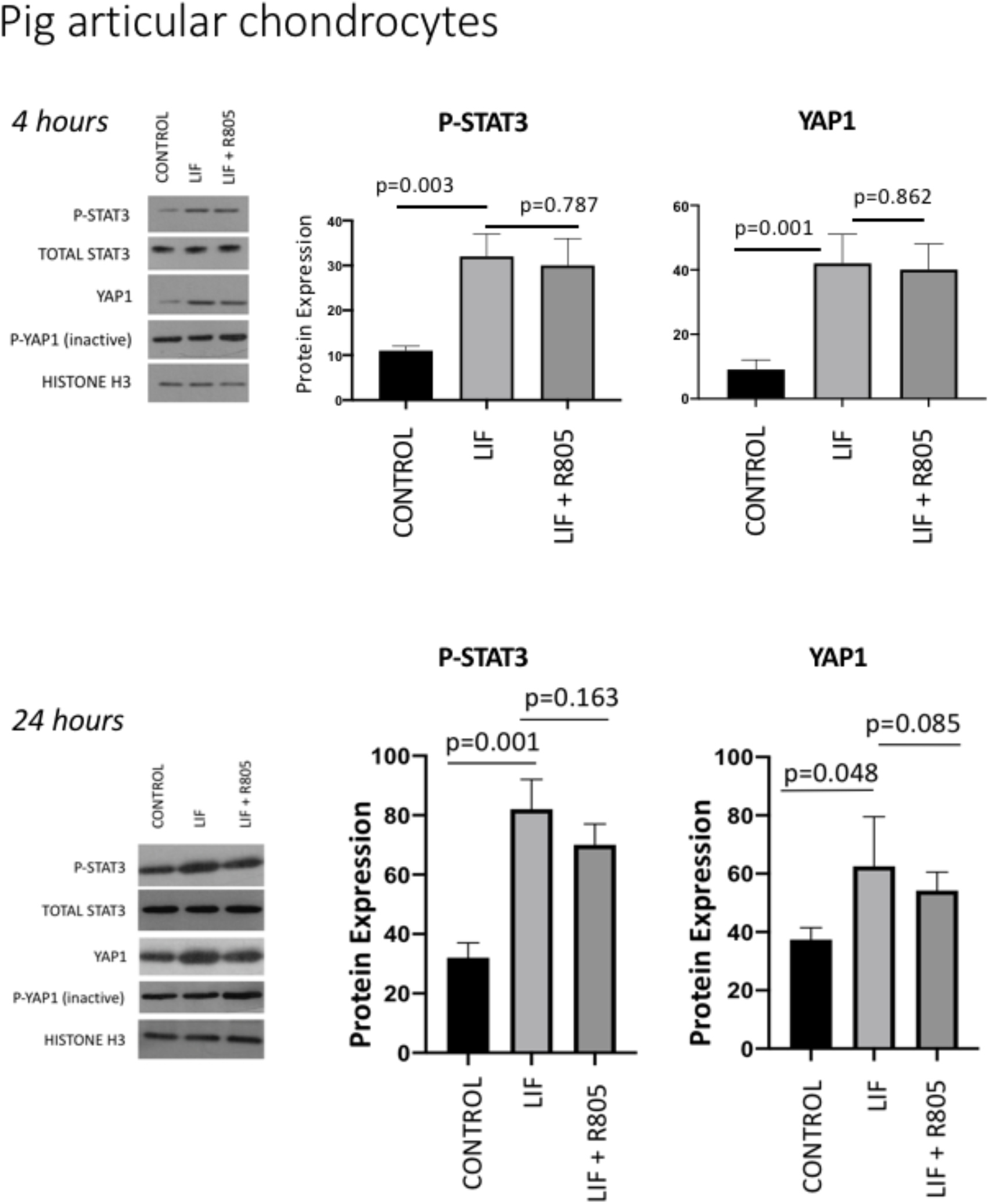
R805 does not affect pro-regenerative signaling downstream of gp130. Protein expression of indicated proteins in pig articular chondrocytes treated with or without LIF and R805 for 4 and 24 hours. Horizontal lines with bars show the mean ± SD. n=3.

**Fig. S20.**
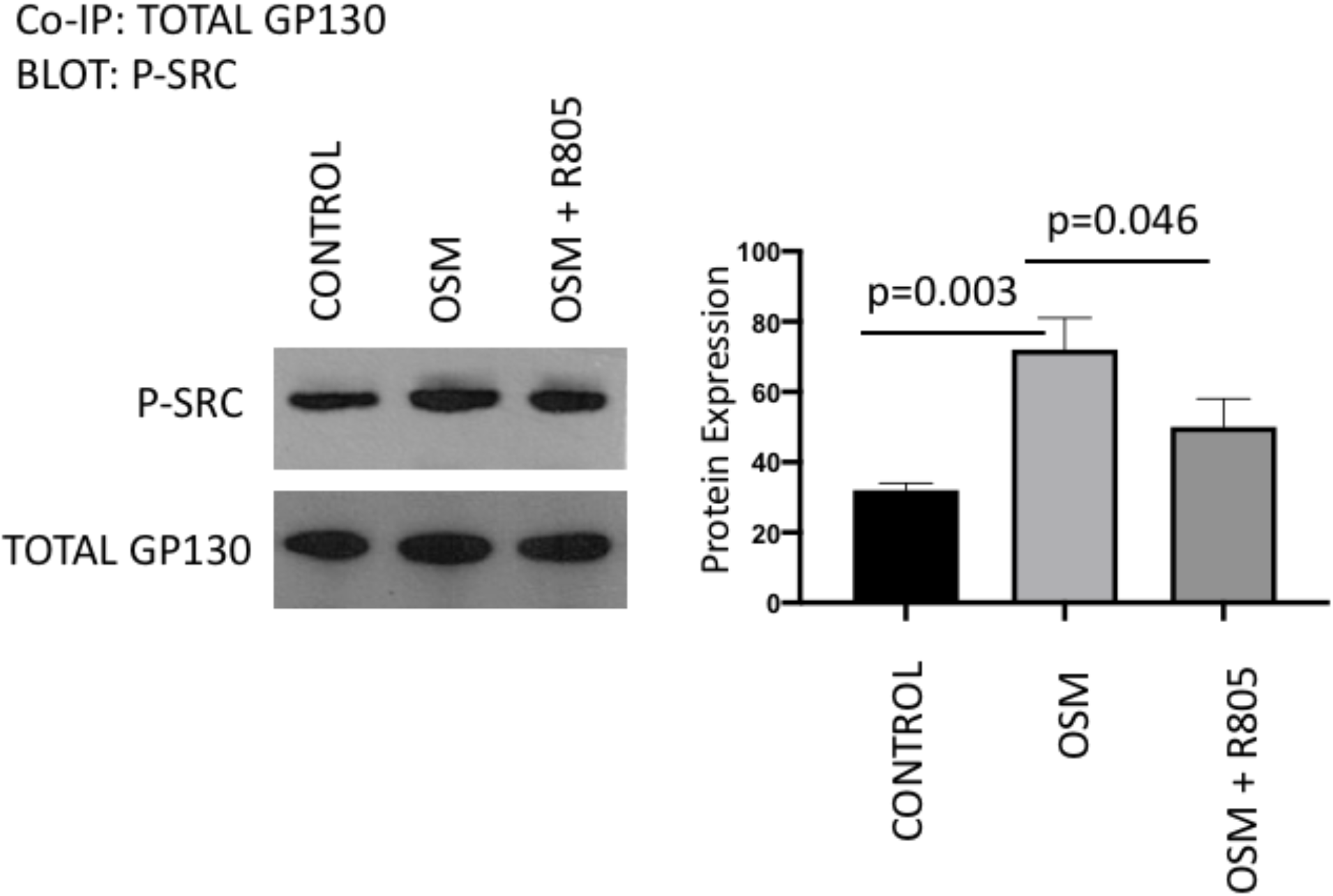
R805 prevents SRC binding to gp130. Levels of complex formation between gp130 and pSRC in pig articular chondrocytes stimulated with or without OSM in presence or absence of R805 for 24 hours. Horizontal lines with bars show the mean ± SD. n=3.

**Fig. S21.**
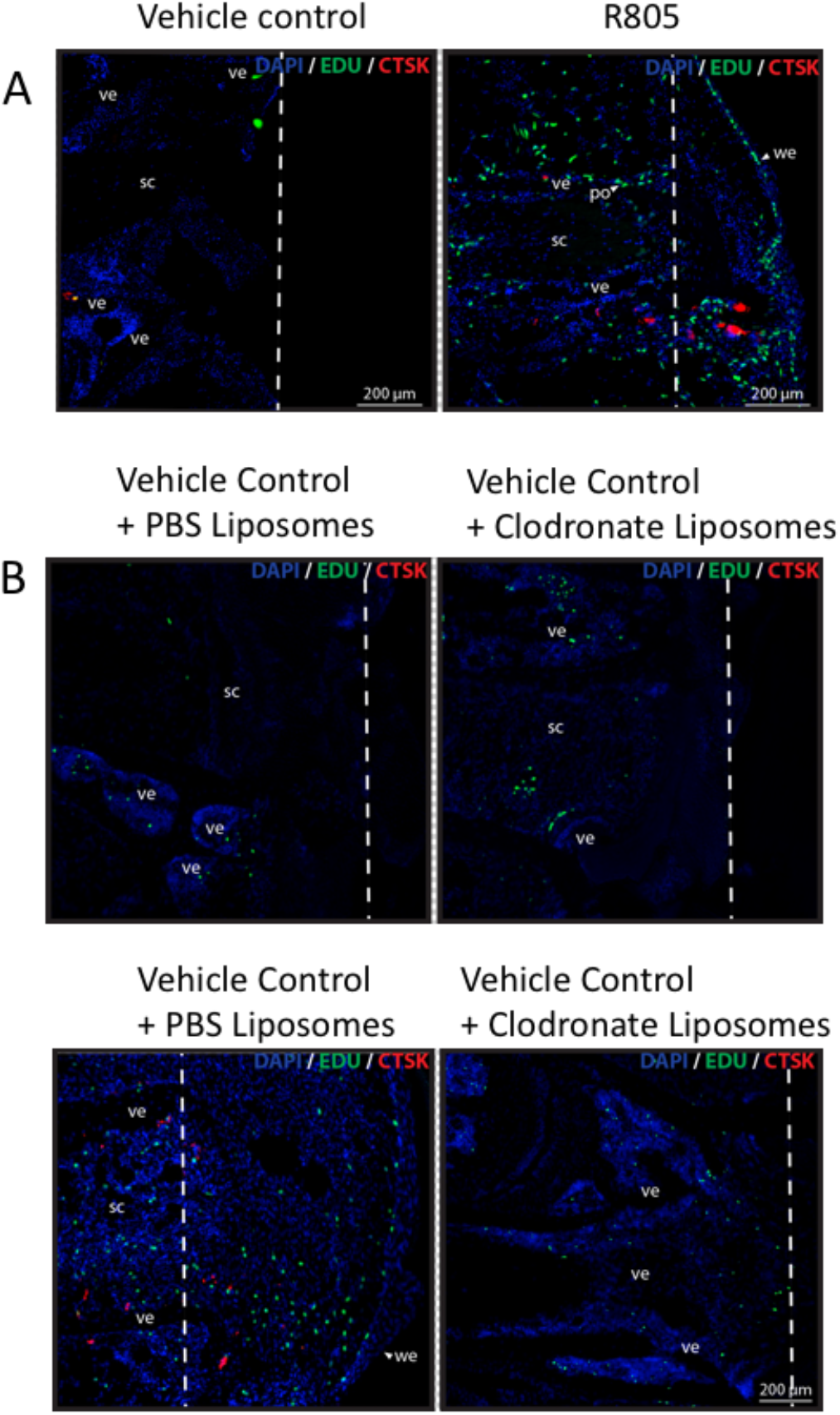
R805 treatment enhances lizard tail regeneration in a macrophage-dependent manner. **A.** EdU and CTSK staining of bearded dragon (*Pogona vitticeps*) tails treated with vehicle control or R805 14 days post-amputation. n=10. po, periosteum; sc, spinal cord; we, wound epithelium; ve, vertebra. **B.** EdU and CTSK staining of bearded dragon (*Pogona vitticeps*) tails treated with vehicle control or R805 14 days post-amputation. n=10. po, periosteum; sc, spinal cord; we, wound epithelium; ve, vertebra.

**Fig. S22.**
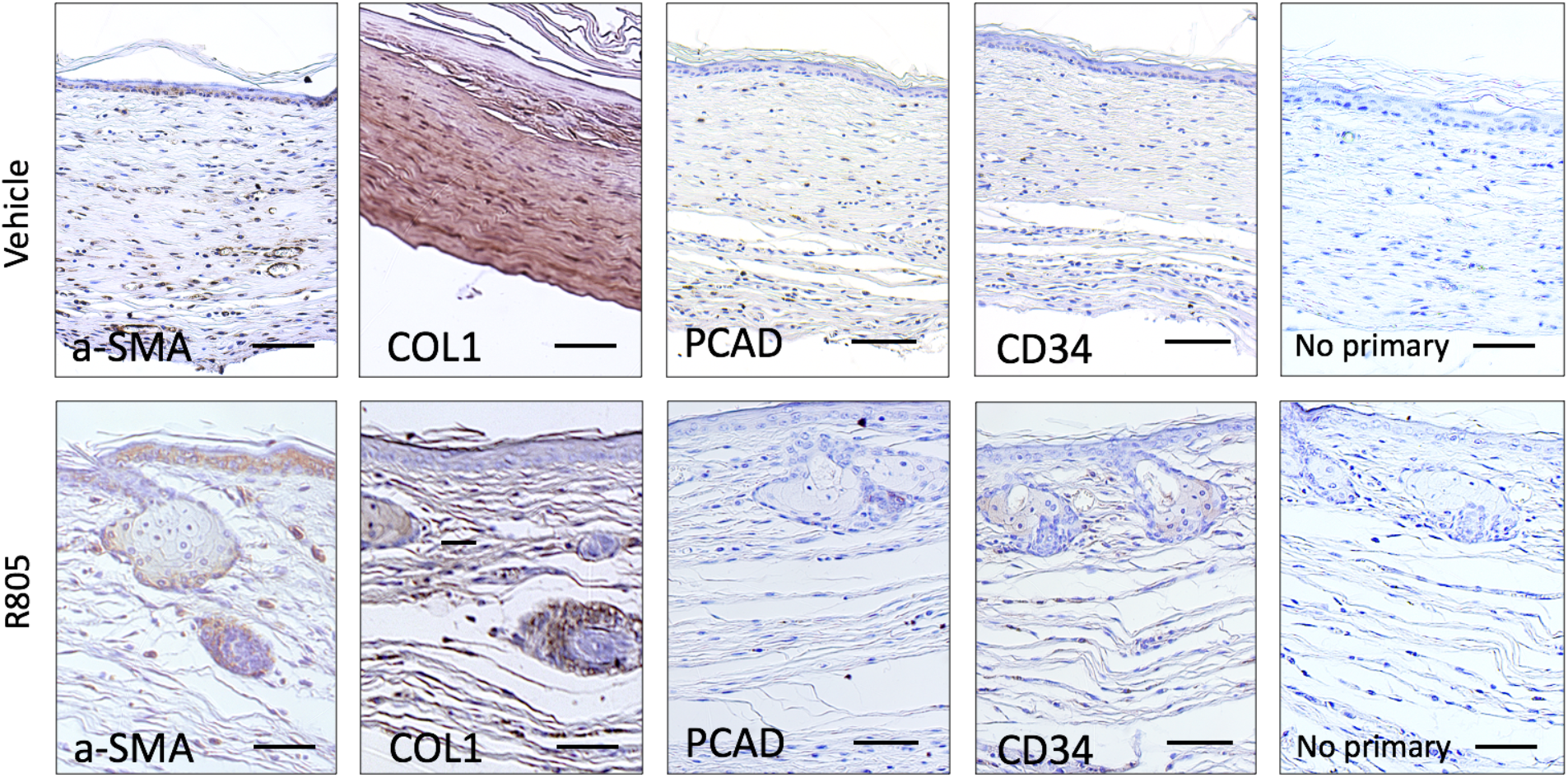
R805-treated mouse exhibits enhanced wound-induced hair neogenesis. Wild type (WT) and R805-treated mouse post-wound day (PWD) 21 wound sections after wound excision. Representative images are shown. n=6.

**Fig. S23.**
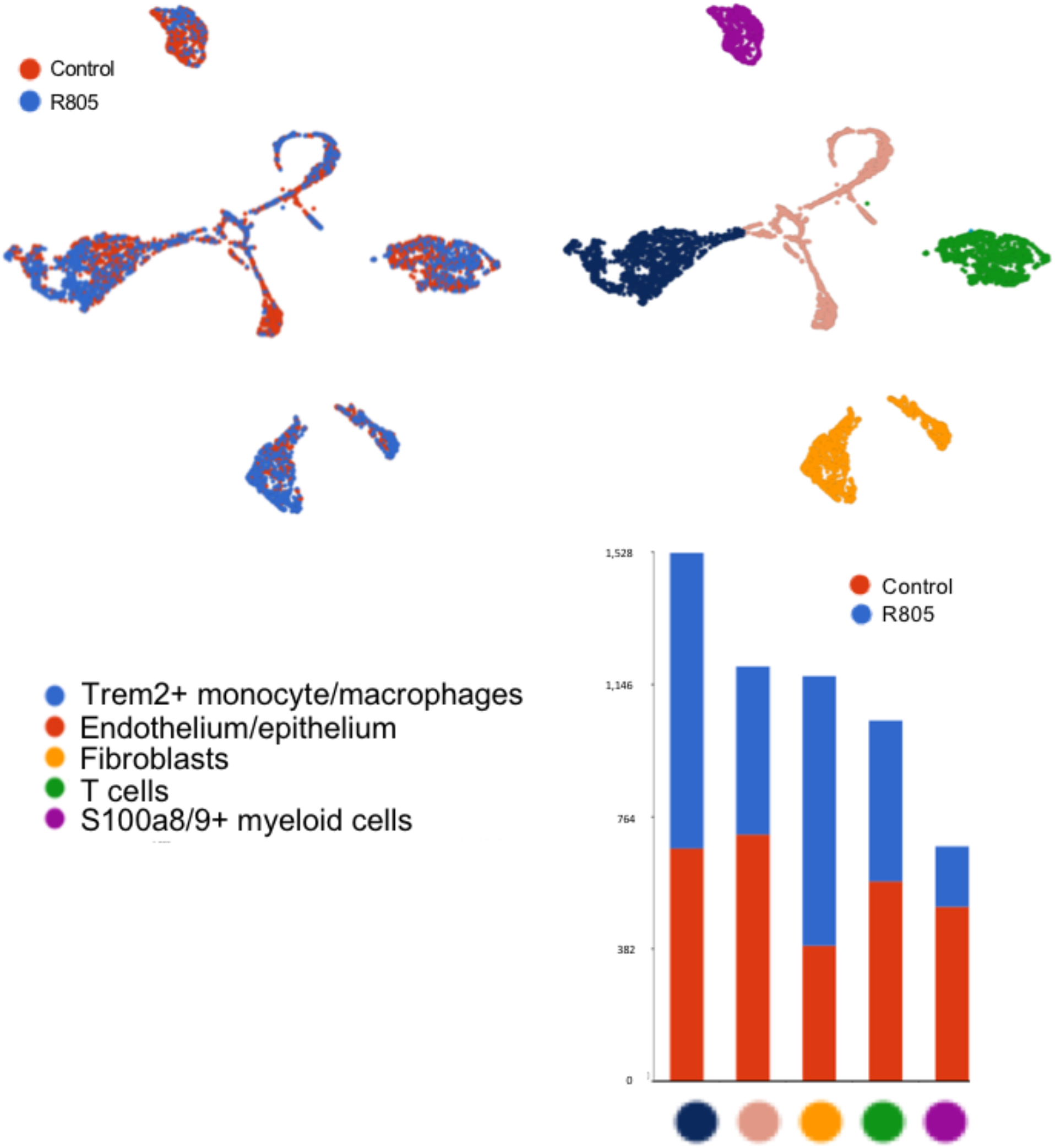
scRNA-sequencing of large wounds from R805-treated and control mice. UMAP and k-means clustering of equal numbers of cells isolated at post wound day (PWD) 14 from the indicated treatment; the number of cells in each cluster from each treatment is displayed in the stacked bar graph. Presumptive identities of each cluster are displayed based on biomarker genes significantly enriched; see also Supp. Table X. Cells were harvested from wounds pooled from 2-3 animals of each treatment and sorted as DAPI and Ter119 negative then subjected to the 10X Genomics workflow.

**Fig. S24.**
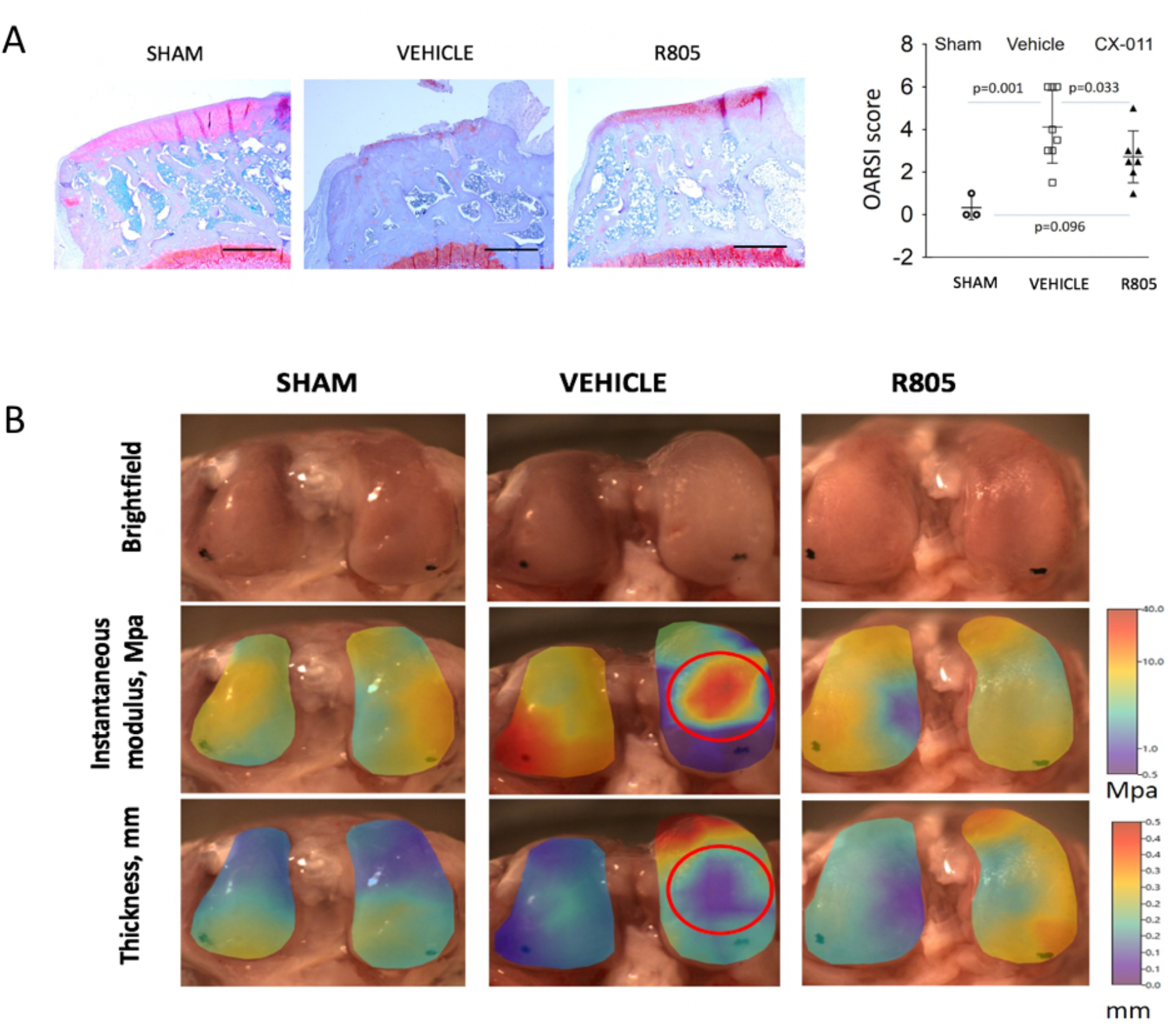
Intra-articular injection of R805 reduces cartilage degeneration following medial meniscal tear surgery in rats. **A.** Intra-articular injection of R805 reduces cartilage degeneration following medial meniscal tear surgery in rats; note increased retention of Safranin O^+^ cartilage (red). Safranin O staining is proportional to the proteoglycan content in the cartilage tissue. Horizontal lines with bars show the mean ± SD. Representative images are shown. n=8. **B.** Automated indentation and biomechanical thickness mapping of ipsilateral femoral condyle of Sham, Vehicle and R805-treated groups. The instantaneous modulus map shows stiff regions (high instantaneous modulus) of the condyle, which suggests bone exposure. R805 maintained thickness and reduced stiffness in rat cartilage. Representative images are shown. n=8. Mpa = megapascals.

**Fig. S25.**
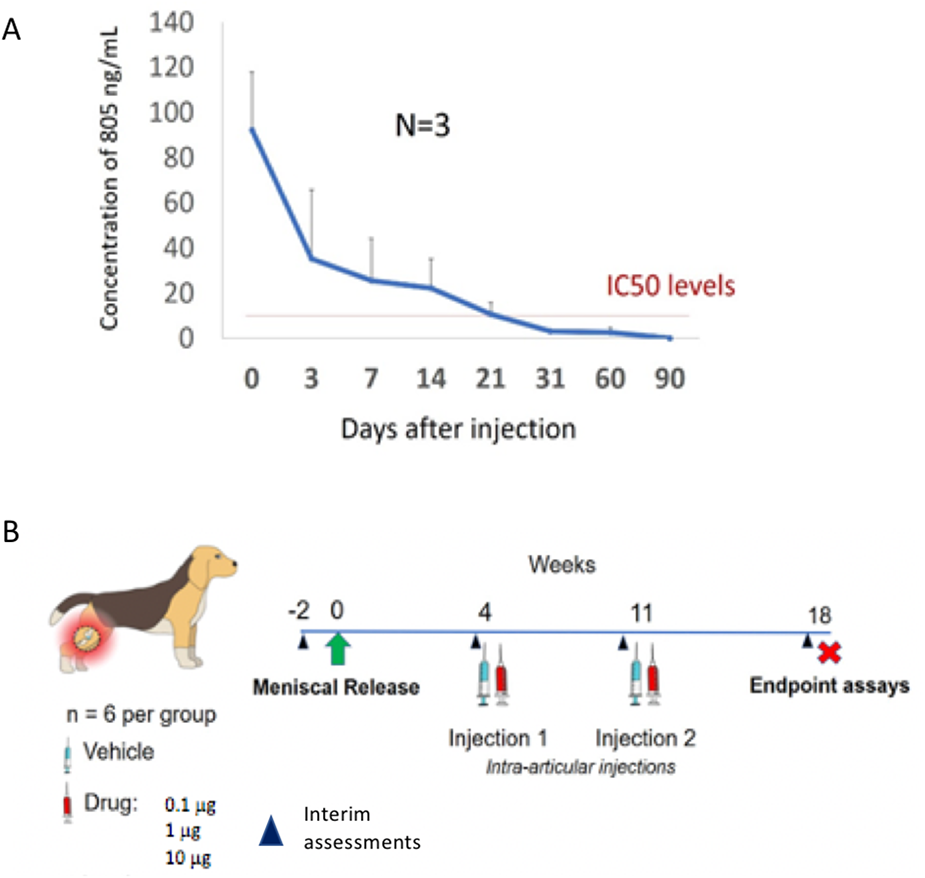
Intra-articular injection of R805 in dog joint. **A.** R805 IC50 concentrations of R805 in the Beagle joint during a 90-day pharmacokinetic study. 1 µg intra-articular injection of R805 was maintained for 3 weeks. The compound was not detected in systemic circulation. n=3. **B.** Schematic depicting the large animal model of osteoarthritis experiment. Intra-articular injections of R805 or vehicle occurred at post-operative weeks 4 and 11. n=7.

**Fig. S26.**
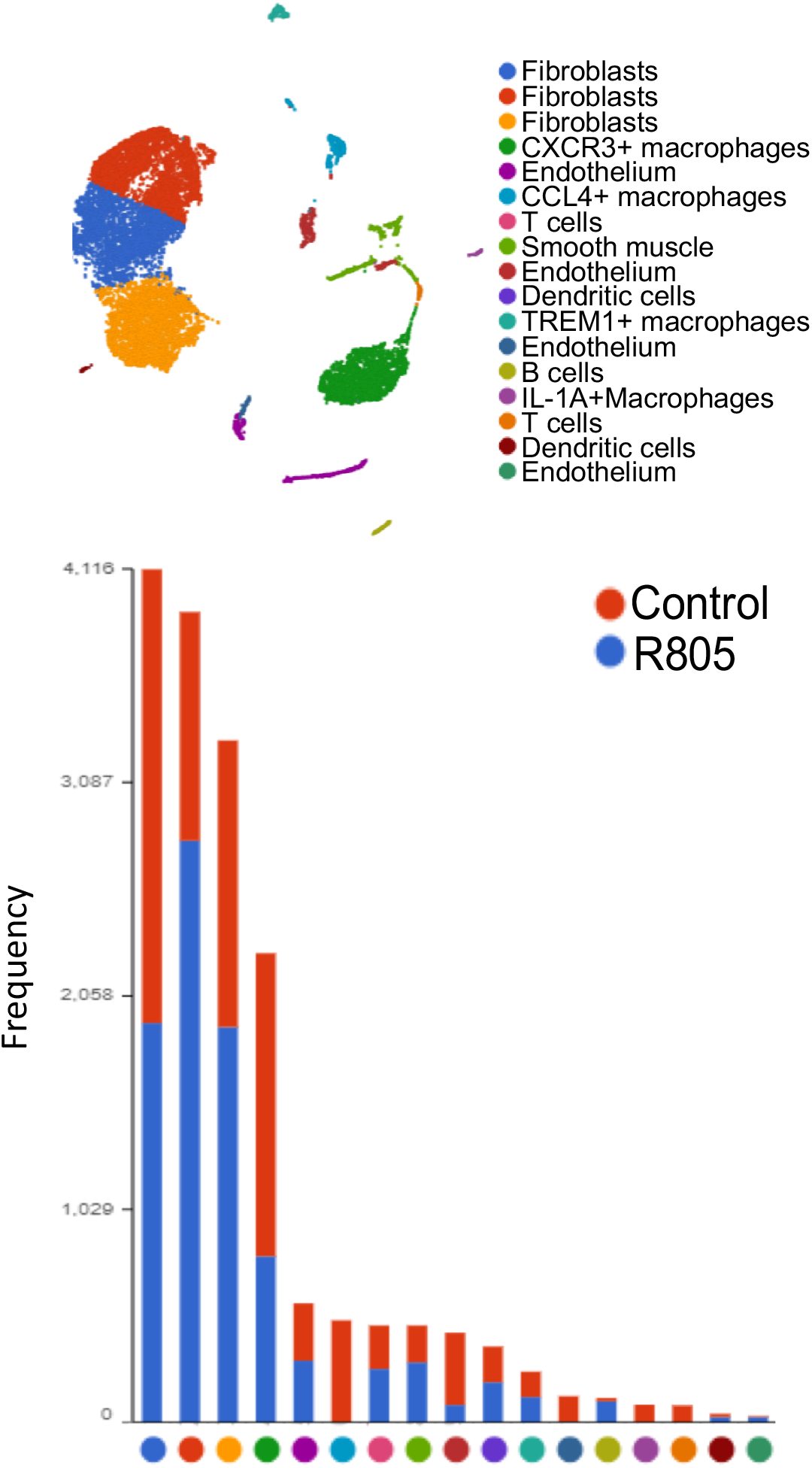
scRNA-sequencing of synoviocytes from R805-treated and control dogs. UMAP and k-means clustering of equal numbers of cells isolated at time of sacrifice from the indicated treatment; the number of cells in each cluster from each treatment is displayed in the stacked bar graph. Presumptive identities of each cluster are displayed based on biomarker genes significantly enriched; see also Supp. Table X. Cells were harvested from synovial membranes pooled from 2-3 animals of each treatment, subjected to RBC lysis, sorted as DAPI negative and then incorporated into the 10X Genomics workflow.

